# Population genetics of bumble bee species with diverging population dynamics

**DOI:** 10.64898/2026.04.14.716867

**Authors:** Asia Piovesan, Christophe Praz, Bernhard Voelkl, Sylvain Lanz, Peter Neumann, Alexis Beaurepaire

## Abstract

Pollinator populations are facing worldwide declines, underscoring conservation needs. Yet, conservation assessments still mostly rely on occurrence data, often derived from heterogeneous and opportunistic observations. While such data can inform on species presence and distribution, they may overlook important markers of population declines. This is particularly problematic for social species such as bumble bees, which typically exhibit low effective population sizes despite high abundance of workers observed in the field. Despite these putative pitfalls, the relationship between occurrence-based and genetic-based estimates remains largely unexplored in social bees. We here investigated spatio-temporal genetic patterns in five Swiss *Bombus* species representing contrasting population trajectories over the last century: *B. humilis* and *B. sylvarum* (stable), *B. ruderatus* (increasing), *B. pomorum* (regionally extinct), and *B. veteranus* (declining). Museum specimens collected between 1929 and 2023 were genotyped at 11 microsatellite loci to compare spatio-temporal fluctuations in genetic diversity and population structure with occurrence data. Overall, multilocus heterozygosity and allelic richness remained stable in all species during the time period investigated, indicating that the diverging population trends did not result in substantial variation of genetic diversity. In contrast, strong and significant shifts in allelic frequencies between time periods were detected in three species, suggesting recent immigration events. Isolation by distance was detected in the cold-adapted *B. veteranus*, while the extant warm-adapted species (*B. humilis*, *B. sylvarum*, *B. ruderatus*) showed high levels of gene flow between locations. In *B. pomorum,* increasing genetic homogenization was observed before extinction. Altogether, these findings show that genetic diversity indexes are not the most adapted tools to monitor conservation status of social bee populations, and that estimates of population structure such as allelic shifts may be more informative. Moreover, these results highlight the importance of monitoring metapopulation dynamics and ensuring connectivity among populations to facilitate gene flow and enable demographic rescue processes.

## 1. Introduction

Pollinators are declining worldwide, compromising both biodiversity and ecosystem services (Díaz et al., 2019; Powney et al., 2019). In Europe, over 30% of bee species are considered near threatened or worse (IUCN, 2025; Michez et al., 2026). Many European bumble bees have undergone population declines since 1900 and are projected to face further contractions under future land use and climate change scenarios, as European environments become increasingly inhospitable (Ghisbain et al., 2024). Yet, the current conservation status of many bee species remains largely unknown (Nieto et al., 2014; Müller and Praz, 2024).

So far, most of the insect conservation efforts and respective legislations rely on species occurrence data, which often come from opportunistic observations rather than standardized monitoring programs. However, occurrence data alone is often insufficient for inferring the conservation status of a species, as it can fail to capture population diversity estimates (Theissinger et al., 2023; Gauthier et al., 2025). This holds particularly true for social Hymenoptera, for which field-based occurrence data can bias conservation statuses estimates due to spatial clustering and other life-history traits. In fact, while workers are most commonly observed the field, usually only queens and males reproduce and genetically contribute to the next generation. Consequently, in social bees the effective population size (Ne, i.e., the number of individuals contributing to the next generation) is always smaller than the census size (Nc). A population might thus appear abundant in field-based data but could be genetically impoverished and be at risk of extinction. As social Hymenoptera such as bumble bees often exhibit low genetic diversity compared to non-social insects (Goulson et al., 2008), this aspect should be taken into account when assessing genetic variation and population dynamics (Webster et al., 2023).

Genetic diversity of a population plays a key role in its adaptation and resilience to environmental stressors and long-term survival. Therefore, there is a growing focus on genetics in species conservation (Beaurepaire et al., 2024; Shaw et al., 2025). Traditionally, conservation studies on bumble bees have relied on estimates of genetic diversity and structure from contemporary populations (Cameron et al., 2011). In addition, the use of historical specimens from entomological collections allows for comparisons of population genetics over time, enabling the identification of genetic fluctuations that may reflect population vulnerability and inform conservation (Leonard, 2008; Lozier et al., 2011).

In this study, museum specimens from five Swiss bumble bee species collected over nearly 100 years were used to determine whether genetic diversity and population structure indexes align with observed population dynamics. In Switzerland 41 bumble bee species have been identified, including both warm- and cold-adapted species (Müller and Praz, 2024). Cold-adapted species are considered more sensitive to heat stress than warm-adapted species (Martinet et al., 2021; Neff et al., 2022). Among the warm-adapted species, *Bombus humilis* and *Bombus sylvarum* show stable occurrence (**Figure S1**). *Bombus ruderatus*, while being common in the past (**Figure S1**), underwent a strong decline from the 1960s until 1990s and started to recover in the late 1990s/early 2000s. The species was classified as “endangered” in 1994 (Amiet, 1994), however, its recent recolonization of the Swiss Midland coupled with increasing population occurrence suggests recent recovery (**Figure S1**). *Bombus pomorum* was widespread in Switzerland until the 1970s but it has not been observed since 1984 and is now considered regionally extinct (RE). Finally, the cold-adapted species *Bombus veteranus* is considered declining due to spatial fragmentation and a temporal decrease in the availability and quality of its habitats leading to its endangered (EN) conservation status.

Using the diverse occurrence trends observed in the five bumble bee species selected, we tested whether genetic diversity indexes match the observed population trends. Additionally, we investigated whether changes in genetic structure over space and time would reveal demographic processes underlying population dynamics, expecting stronger shifts in genetic structure in species exhibiting the most pronounced population fluctuations. Given the noticeable temperature increase observed in Switzerland in the past decades and its potential consequences on species diversity and distribution (Vittoz et al., 2013), we compared extant warm- and cold-adapted species and hypothesized to identify signs of genetic vulnerability in the cold-adapted species due to their decreasing habitat ranges and heat stress vulnerability while expecting maintained gene flow in warm-adapted species.

## 2. Material and methods

### 2.1 Species selection

In total, four warm-adapted species (*Bombus humilis, B. sylvarum, B. ruderatus,* and *B. pomorum*) and one cold-adapted species (*Bombus veteranus*) displaying different population dynamics in Switzerland from 1900 to 2023 (**Figure S1**) were selected. The full list of specimens was extracted from a 1 x 1 km square resolution database provided by the Swiss data center Info Fauna (www.infofauna.ch) (**Table S1**). Species trends were based on examination of occurrences over time (Figure S1) obtained from Müller and Praz (2024).

### 2.2 Study sites

For each species, diploid females sampled between 1929 and 2023 in four Swiss cantons (Geneva (GE), Vaud (VD), Neuchatel (NE) and Bern (BE)) were selected (**Figure 1**). These cantons were chosen as all five species occur there and the coverage in specimens is continuous over the examined time periods. Only specimens collected below 1000 m elevation were selected. For *B. veteranus*, samples from other cantons were included due to the scarcity of specimens available in collections for the selected areas. To minimize the likelihood of sampling nest siblings, only one specimen per 1 x 1 km square location was selected for each sampling year in most cases. When sample size was too limited to do this, relatedness was analyzed using the software COLONY (Jones and Wang, 2010) and when a sibship probability of more than 0.95 was found, one individual was removed. Based on these criteria, one *B. veteranus* specimen was excluded (**Table S1**).

**Figure 1.**
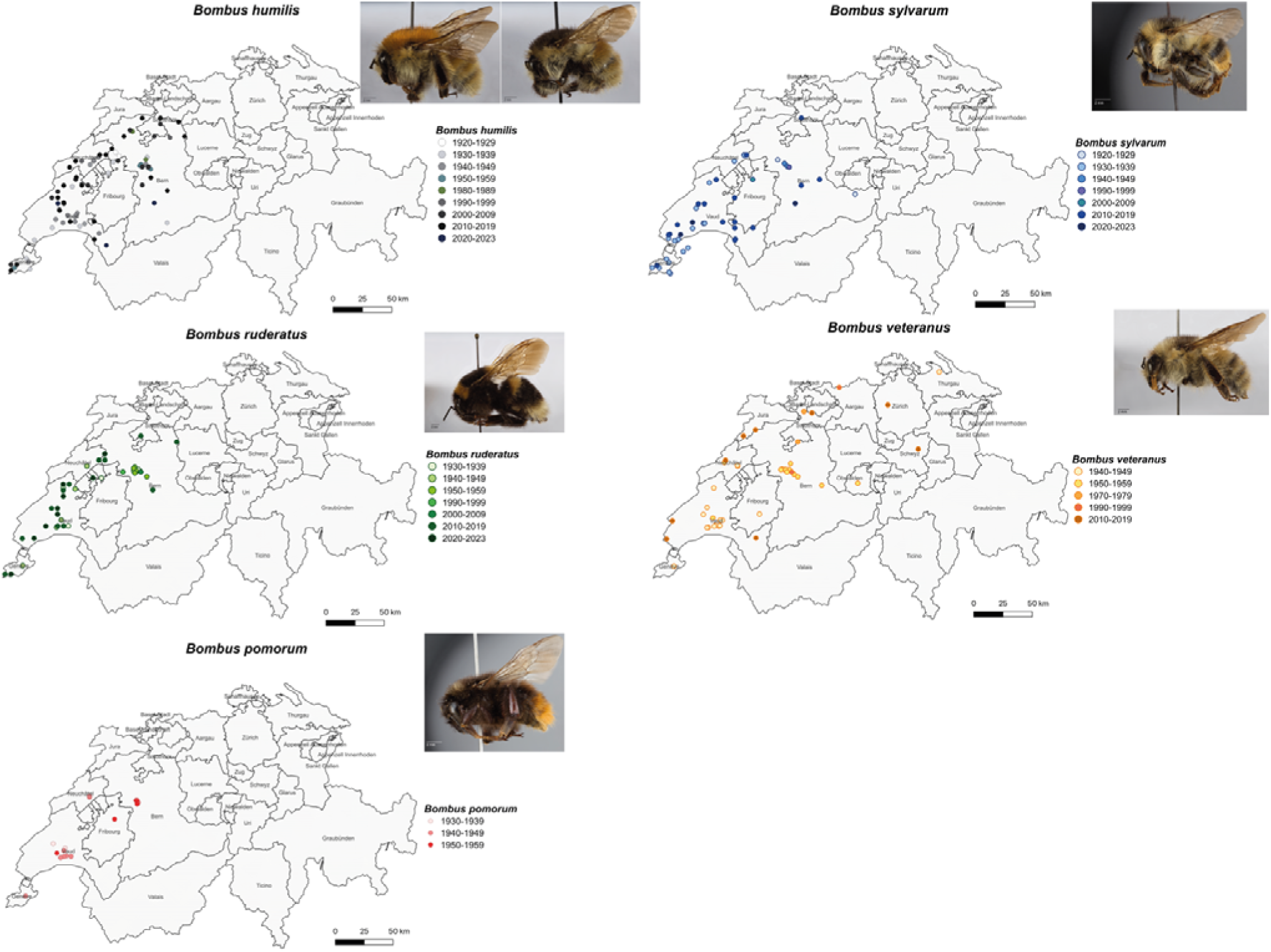
Sampling time and location of the specimens genotyped. For each species, the genotyped specimens were grouped by decade. Pictures taken using a Canon EOS 6D camera and processed with the programs Adobe Photoshop Lightroom and Helicon focus v5.3 illustrate specimens housed at the Natural History Museum of Lausanne.

### 2.3 DNA acquisition, extraction and amplification

In total, 435 female specimens from the Swiss Hymenoptera collections of *Bombus humilis* (N=123), *Bombus ruderatus* (N=87), *Bombus veteranus* (N=79), *Bombus sylvarum* (N=94) and *Bombus pomorum* (N=52) sampled from 1929 to 2023 were selected. One mid leg was sampled from each specimen and DNA was extracted using NucleoSpin® Tissue extraction kit (Macherey-Nagel, Oensingen, Switzerland). Eleven microsatellite loci (Estoup et al., 1996; Reber-Funk et al., 2006) initially developed for *B. terrestris* and *B. lucorum* and successfully used to genotype museum specimens of European bumblebees in Maebe et al. (2016) were targeted (**Table S2**). Single PCR reactions were performed in 10 μL using 2 μL of extracted DNA, 0.3 μL for each forward and reverse primer, 2 μL Taq Buffer, 0.05 μL of Taq Polymerase (Labgene.ch) and MilliQ water as follows: initial denaturation for 5 min at 94 °C, 35 cycles of denaturing at 92°C for 30 sec, annealing at the correspondent temperature of each primer (**Table S2**) for 30 sec, elongation at 72 °C for 45 sec, and the final extension for 10 min at 72 °C. Microcapillary electrophoresis of the final PCR products was carried out by Microsynth AG (Balgach, Switzerland) and the genotypes were manually scored using the Peak Scanner software v1.0 (Applied Biosystems ®). A genotyping threshold of 50% was applied, retaining only individuals successfully genotyped at 50% or more of the loci for genetic analyses. To maximize the study resolution and detect fine scale genetic variations, changes in genetics over space and time were assessed at the individual level (Milligan et al., 2018; Rohde et al., 2024).

### 2.4 Genetic analyses

All analyses listed below were conducted separately for each species, with marker sets varying among species due to the exclusion of monomorphic markers (**Table S3**).

The software ADZE (Szpiech et al., 2008) was used to calculate rarefied allelic richness to assess whether the total sample size of each species sufficiently represented their genetic diversity. Hardy-Weinberg assumptions were tested with GENEPOP v.4.7.5 (Raymond and Rousset, 1995) using the Markov chain Monte Carlo (MCMC) method with 1000 dememorization steps, 100 batches and 1000 interactions per batch. Linkage disequilibrium between pairs of markers overall specimens was tested using the same software. Observed heterozygosity per locus was calculated using Genalex 6 (Peakall and Smouse, 2006).

#### 2.4.1 Population genetic diversity

For each specimen multilocus heterozygosity was calculated using the *MLH()* function in the R v4.3.2 (R Core Team, 2023) package *inbreedR* (Stoffel et al., 2016). Due to the heteroscedasticity of heterozygosity data, a robust linear regression model was fitted using the function *lm_robust()* from the R package *estimatr* (Blair et al., 2018) to assess the relationship between multilocus heterozygosity and time, modelled as *(multilocus heterozygosity ∼ time)*. In *B. ruderatus*, multilocus heterozygosity was calculated for specimens collected before (1930-1960) and after (1990-2023) the hypothesized population decline before recovery, as well as for the entire time period (1930-2023).

Next, temporal changes in allelic occurrence was investigated using a Bayesian multilevel logistic regression. Each observation *y_ijkl_* indicated whether allele *l* at locus *k* was detected in individual *i* sampled in year *t*.

The model was specified as: *y_ijkl_∼Bernoulli(p_ijkl_), logit(p_ijkl_) = β_0_ + β_1_ year_scaled_t_ + u_k_* + *v_kl_* + *w_i_*, where β_0_ is the intercept and β_1_ is the fixed effect of sampling year (scaled to mean 0 and unit variance). Random intercepts were included for locus (*u_k_∼N (0, σ^2^_locus_)*), allele nested within locus (*v_kl_∼N (0, σ^2^_allele_)*), and individual (*w_i_∼N (0, σ^2^_individual_)*). This hierarchical structure accounts for differences in baseline allele frequency among loci and alleles and for the non-independence of multiple allele observations originating from the same individual. The coefficient *β_1_* therefore quantifies the average temporal change in the probability that an allele variant is observed in a sampled individual, after accounting for locus-, allele-, and individual-level heterogeneity.

The model was fit in a Bayesian framework using Stan (Stan Development Team, 2025) via the R package *brms* (Bürkner, 2017) with 4 Markov chains of 6000 iterations each. Convergence and model adequacy were assessed using standard posterior diagnostics and posterior predictive checks.

Specimens collected the same year were then grouped into temporal clusters (**Table S1**) to estimate inbreeding coefficients (Fis) with Fstat (Goudet, 1995). *B. veteranus* and *B. pomorum* were excluded from the analyses as too few data points were available, and confidence intervals were too wide to allow reliable estimation. To investigate temporal trends in inbreeding, a linear regression model was fitted in R v4.3.2 for Fis as functions of sampling years (time): *Fis ∼ time*. In *B. ruderatus*, the inbreeding coefficient was calculated for specimens collected before (1930-1960) and after (1990-2023) the hypothesized population decline before recovery as well as for the entire time period (1930-2023). Model residuals were visualized with Quantile-Quantile plots (QQ plots) to assess adherence to normality assumptions.

#### 2.4.2 Population genetic structure

A Principal Component Analysis (PCA) was performed with the R packages *adegenet* (Jombart, 2008) and *ellipse* (Murdoch and Chow, 2020). To assess the association between spatial and temporal distances and genetic variation, the average temporal and spatial distances of each specimen to all others were calculated and scaled (mean=0; SD=1). Pearson correlation between the scaled temporal distances and spatial distances and the first two principal components (PC1 and PC2) of the PCA were then calculated. To quantify the extent to which spatial and temporal predictors explain multivariate genetic variation, a redundancy analysis (RDA) was performed with the R package *vegan* (Dixon, 2003) using the function *rda()*. The multivariate allele presence-absence matrix was used as response variable *y*, while sampling year and coordinates (latitude and longitude) were included as explanatory variables. All predictors were centered and scaled (mean=0; SD=1). First, a full model was fitted as *y ∼ scale(year) + scale(longitude) + scale(latitude)* to quantify the total proportion of genetic variation explained jointly by time and space. To disentangle the independent contributions of spatial and temporal predictors, partial RDAs were performed by fitting respectively *y ∼ scale(year) + Condition(scale(longitude) + scale(latitude))* and *y ∼ scale(longitude) + scale(latitude) + Condition(scale(year)).* Models’ significance was assessed using permutation-based ANOVA (999 permutations) and marginal effects of individual predictors were evaluated using permutation tests with terms assessed sequentially. The proportion of explained variance was quantified using adjusted R² values.

The relationship between genetic differentiation and spatiotemporal factors was investigated through linear mixed-effects models. Pairwise genetic distances based on the PhiPT distance index were calculated in Genalex 6 (Peakall and Smouse, 2006), pairwise geographic distances were calculated as Euclidean distances between GPS coordinates, and pairwise temporal distances were measured as the absolute difference in collection years. All matrices were standardized using z-score normalization by subtracting the mean and dividing by the standard deviation of all pairwise values (excluding missing values), resulting in matrices with mean 0 and standard deviation 1. The outer correlation between the temporal and geographical distances was calculated as the Pearson correlation. A linear mixed-effects model (function *lmer* from the R package *lme4* (Bates et al., 2003)) was formulated as: *genetic_distance ∼ geographic_distance * temporal_distance + (1 | id1) + (1 | id2)*, were id1 and id2 represent the individuals of each pair. Random intercepts for id1 and id2 were included to account for non-independence due to repeated appearances of individuals in pairwise comparisons. Finally, the significance of fixed effects was assessed based on Satterthwaite’s method for estimating degrees of freedom using the R package *lmerTest* (Kuznetsova et al., 2013). The effect of spatial and temporal distances on genetic distance was plotted with *ggplot2* package (Wickham, 2011). The residuals distribution was then plotted in a QQ plot. Model fit was further evaluated using marginal Rand conditional R² from the R package *performance* (Lüdecke et al., 2021). Model assumptions were checked via residual diagnostics and QQ plots.

## 3. Results

In total, 366 specimens were successfully amplified at ≥ 50% loci, including 104 *Bombus humilis*, 81 *Bombus ruderatus*, 59 *Bombus veteranus*, 80 *Bombus sylvarum* and 42 *Bombus pomorum* (**Table S1**). According to the rarefaction analysis (**Figure S2**), the curves reached a plateau with 50-70 individuals per species, suggesting that sufficient samples were used in most of the species.

Deviations from HWE were detected in several species and time periods, but not consistently across decades or species (**Table S4**). Pairwise linkage disequilibrium analyses revealed that only a few pairs of loci showed significant deviations, suggesting partial linkage effects between markers (**Table S5**). Most loci exhibited high genetic variability, with observed heterozygosity values greater than 0.5, yet a few markers showed low levels of polymorphism (**Table S6**). Deviations from linkage disequilibrium were species-specific and mostly involved loci with low genetic diversity. Because these deviations were not consistent among species and reflected low marker variability, all markers were retained in subsequent analyses.

### 3.1 Temporal patterns of populations genetic diversity

The analysis of multilocus heterozygosity over time revealed contrasting trends between genetic diversity and demographic changes across species (**Figure 2**, **Table S7**).

**Figure 2.**
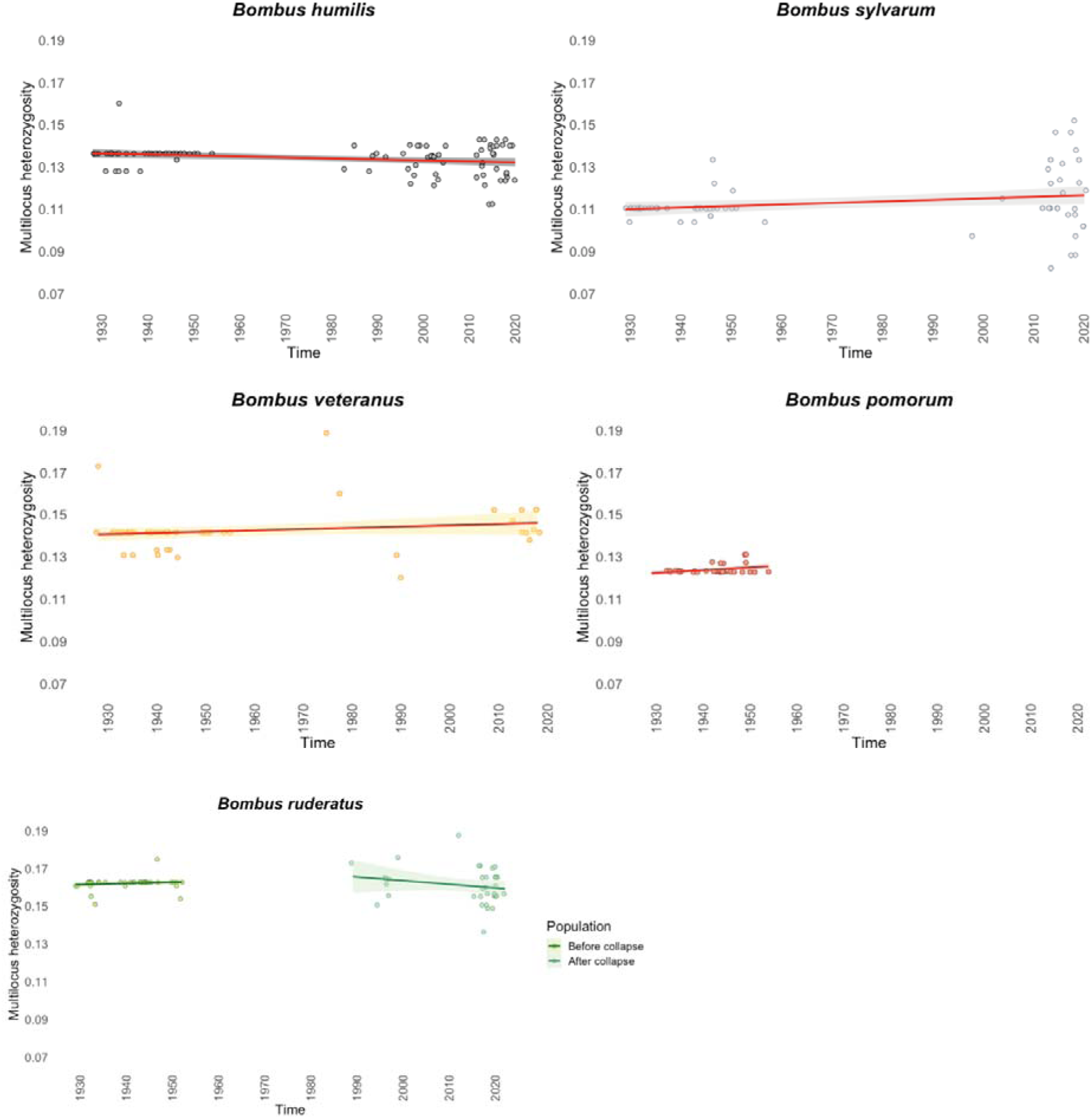
Temporal fluctuations in multilocus heterozygosity in each bumble bee species. The line indicates the estimate of a robust linear regression; the shaded area gives the 95% C.I. for the regression line. For *B. ruderatus* linear relations between multilocus heterozygosity and time have been estimated separately for the periods before (1930-1960) and after (1990-2023).

The stable species *B. humilis* displayed a significant but low decrease of heterozygosity (slope = −4.64×10^-05^, *p* = 1.86×10^-02^) while *B. sylvarum* showed stable heterozygosity over time (*p* = 0.085). Overall, the heterozygosity levels of *B. ruderatus* remained stable through time (*p* = 0.954) (**Figure S3**). No significant changes in heterozygosity occurred before *B. ruderatus* collapse (*p* = 0.417) as well as during its recolonization phase (*p* = 9.82×10-02). The declining species *B. veteranus* exhibited stable heterozygosity over time (*p* = 0.123). A significant increase in heterozygosity was measured in *B. pomorum* before its collapse in 1980s (slope: 0.0001, *p* = 0.03) (**Figure 2**, **Table S7**). Across all species, adjusted R² values varied between 0 and 0.131, indicating that the models had modest explanatory power and that the temporal trends accounted for a small portion of heterozygosity variation.

No strong or consistent temporal fluctuations in allele richness was observed (**Figure 3**, **Table S8**) and all species exhibited allelic variation. All models converged well indicating reliable posterior estimates (**Table S8**) and posterior predictive checks showed no evidence of model misspecification (**Figure S4**). No significant changes in Fis were observed in *B. humilis* (*p* = 0.147), *B. sylvarum* (*p* = 0.589) and *B. ruderatus* (*p* = 0.867) (**Figure 4**, **Figure S5**, **Table S9**). Residuals followed a normal distribution (**Figure S6**).

**Figure 3.**
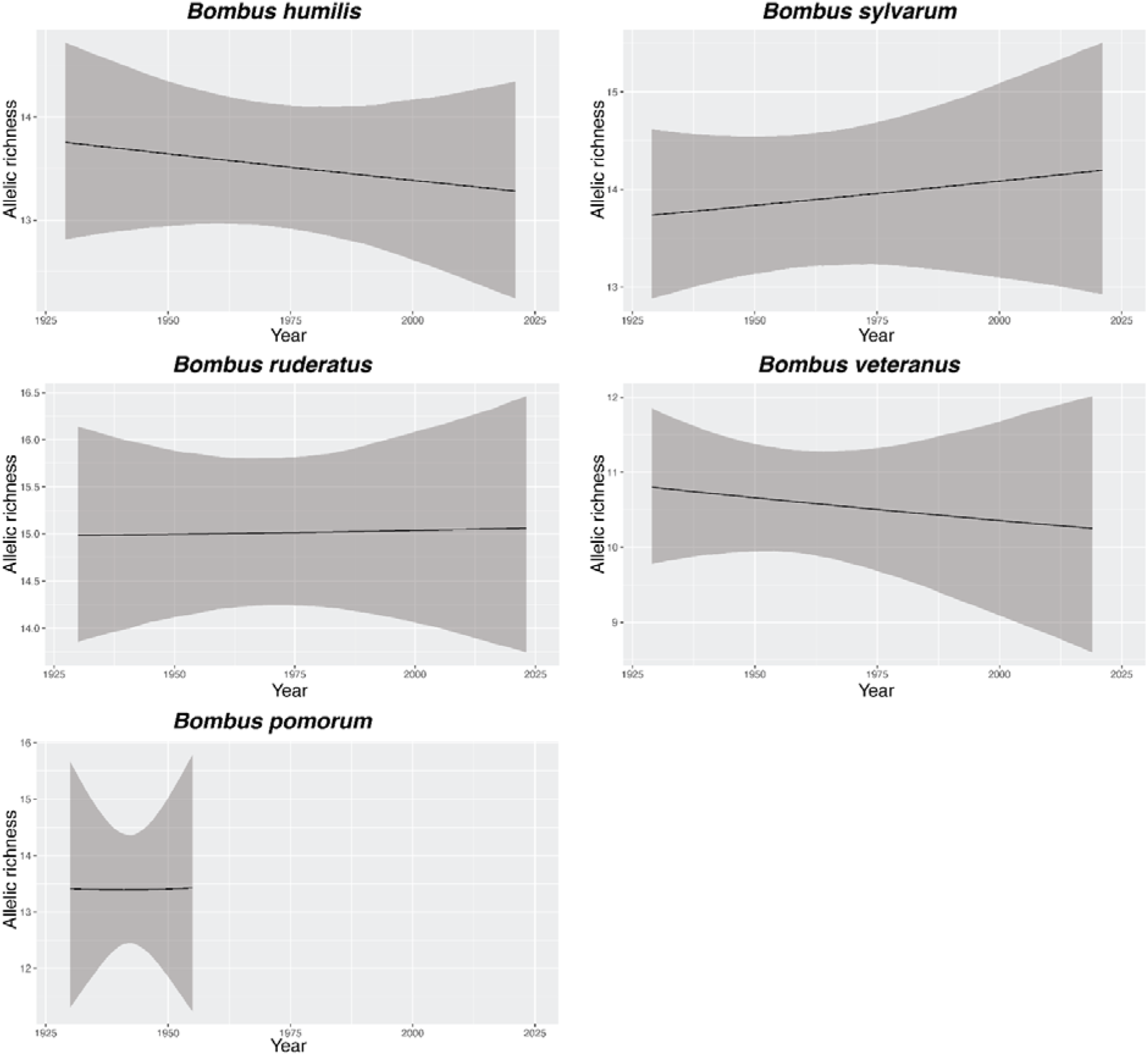
Predicted change in allelic occurrence over time for each species. The solid line indicates the posterior mean prediction obtained from Bayesian logistic mixed-effects models. The shaded area shows the 95% Bayesian credible interval of the predicted probability of allele presence. Model fit based on four Markov chains, and 6000 iterations per chain.

**Figure 4.**
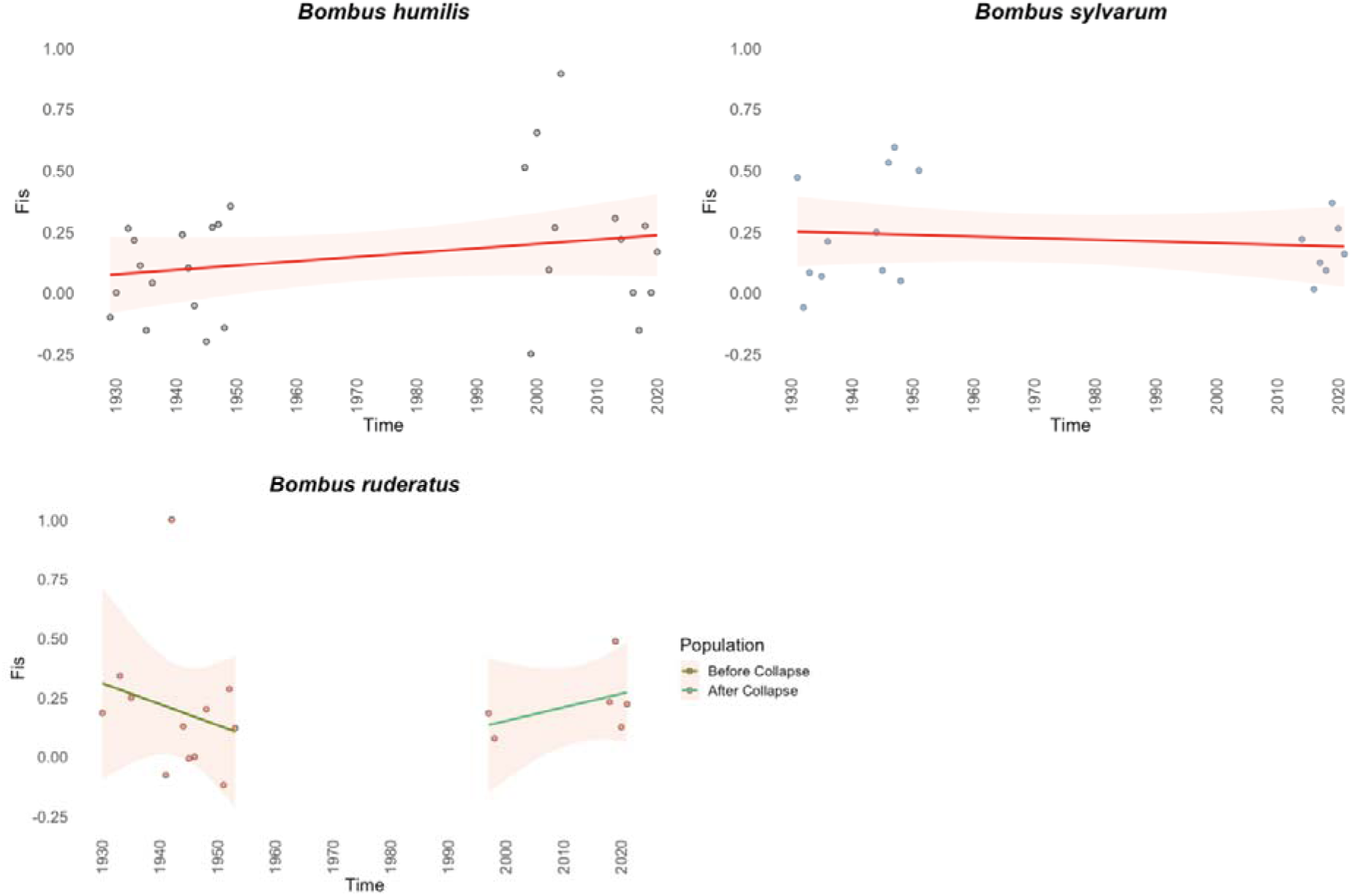
Inbreeding coefficient (Fis) of *B. humilis*, *B. sylvarum* and *B. ruderatus* from 1930 to 2023 in Switzerland. Development of inbreeding (Fis) over time of *B. humilis*, *B. sylvarum* and *B. ruderatus*. The line indicates the estimate of a linear regression; the shaded area gives the 95% C.I. for the regression line. For *B. ruderatus* linear relations between inbreeding and time have been estimated separately for the periods before (1930-1960) and after (1990-2023).

### 3.2 Spatio-temporal population structure

#### 3.2.1 Principal Component Analysis (PCA)

*B. humilis* showed a gradual shift in allelic composition over time (**Figure 5**). A significant negative correlation between temporal distance with PC1 (*r* = –0.279, *t* = –2.94, *p* = 0.004) was detected, while the correlation with PC2 was not significant. Spatial distance was not significantly correlated with either PC1 or PC2 (**Table S10**). *B. sylvarum* individuals grouped together forming a single cluster, suggesting no temporal pattern (**Figure 5**). No significant correlations were found for both temporal and spatial distance with either PC1 or PC2 (**Table S10**). *B. ruderatus*, which went through a recent recolonization after population collapse before 1990s, displayed two distinct genetic clusters (**Figure 5**). Temporal distance showed a significant positive correlation with PC1 (*r* = 0.796, *t* = 11.69, *p* < 2.2×10^-16^) and a significant negative correlation with PC2 (*r* = –0.250, *t* = –2.30, *p* = 0.024). Spatial distance showed no significant association with either PC1 or PC2 (**Table S10**).

**Figure 5.**
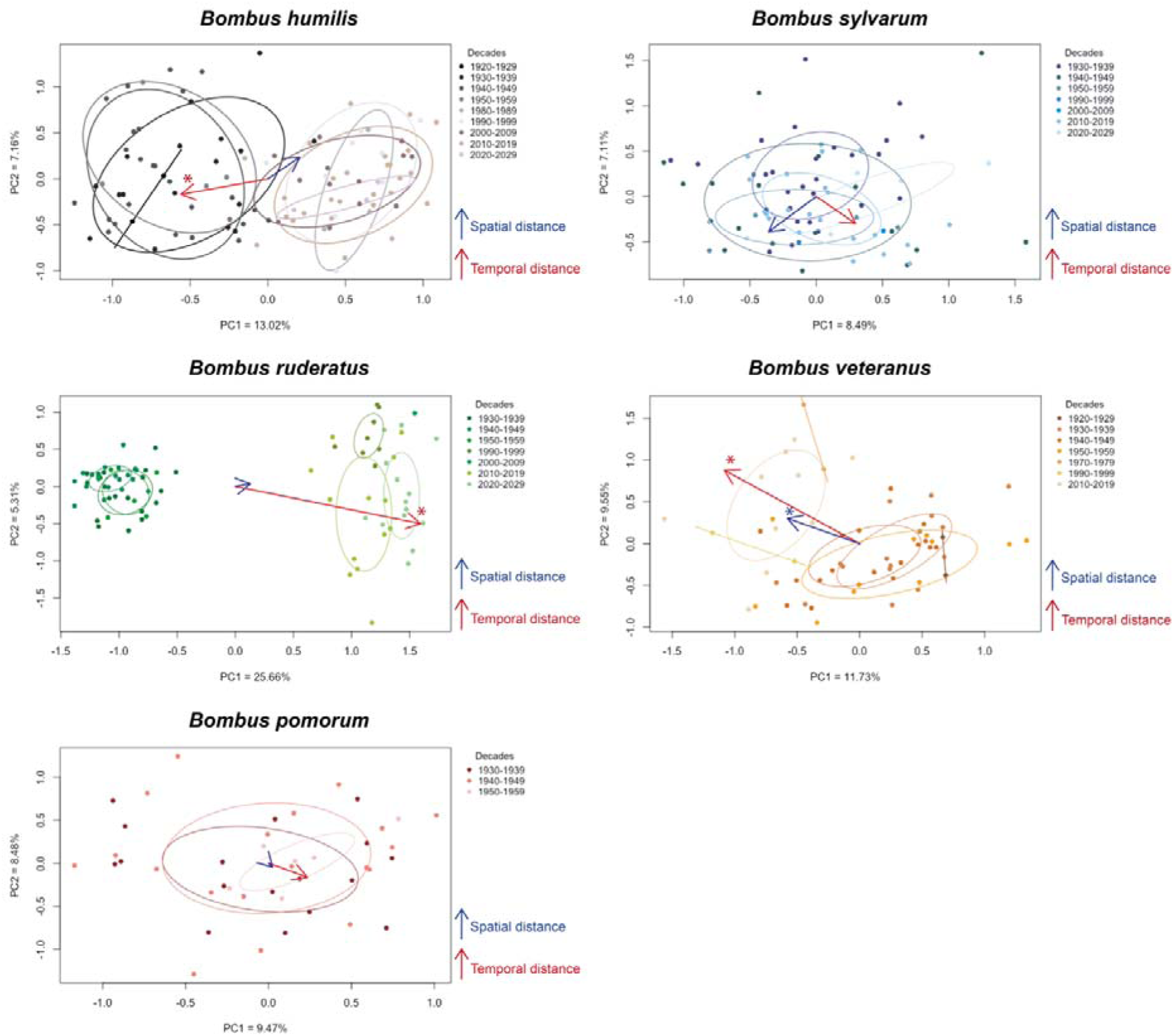
Principal Component Analysis (PCA) based on microsatellite data of each bumble bee species. The plot is displayed according to the two first Principal Components (x-axis: PC1 and y-axis: PC2) and the percentage of variability represented by each component is also reported. Genotype distribution according to time was done by assigning each specimen to the corresponding decade. For each decade with at least two individuals, confidence ellipses based on the covariance of PCA were drew. Arrows represent the correlations between the first two principal components (PC1 and PC2) and spatial (blue) or temporal (red) distances, with their length and direction proportional to the strength and sign of the correlation. Significant correlations (*p* < 0.05) are indicated with a *.

The declining species *B. veteranus* showed no apparent clustering (**Figure 5**). Temporal distance showed a significant negative correlation with PC1 (*r* = –0.542, *t* = –4.91, *p* = 7.86×10^-16^) and a significant positive correlation with PC2 (*r* = 0.439, *t* = 3.72, *p* = 0.0005). Spatial distance exhibited significant negative correlation with PC1 (*r* = –0.289, *t* = –2.30, *p* = 0.025), but not with PC2 (**Table S10**). Finally, in the currently extinct species *B. pomorum* a single cluster and no temporal pattern was identified. Both temporal and spatial distance were not significantly associated with PC1 or PC2 (**Table S10**).

#### 3.2.2 Redundancy analysis (RDA)

An RDA was conducted to quantify the extent to which spatial and temporal variables explain patterns of genetic variation. In *B. humilis*, the combined effect of space and time explained 9.2% of the total genetic variance. Sequential testing of individual predictors indicated that time had a significant effect (F = 11.43, *p* = 0.001), while longitude and latitude had not. Partial RDA showed that time alone explained 8.9% of the variance (F = 10.86, *p* = 0.001) while space (∼0.01%) was had no significant effect (**Table S11**). In *B. sylvarum*, the combined effects of space and time explained 4.2% of the total genetic variance. Sequential tests indicated that time contributed significantly (F = 2.13, *p* = 0.001), while longitude and latitude did not. Partial RDA indicated that time explained 2.6% of the variance (F = 2.08, *p* = 0.001) while space (∼1.5%) had no significant effect (**Table S11**). In *B. ruderatus*, the combined effects of space and time significantly explained 26.6% of the total genetic variance. Sequential tests indicated that time contributed significantly (F = 24.77, *p* = 0.001), while longitude and latitude were not significant. Partial RDA revealed that time alone significantly explained 23.3% of the total variance (F = 24.80, *p* = 0.001) while space significantly explained 1.0% (F = 1.54, *p* = 0.001) (**Table S11**). In *B. veteranus*, the combined effects of space and time explained 12.3% of the total genetic variance. Sequential tests indicated that year (F = 4.58, *p* = 0.001), longitude (F = 1.56, *p* = 0.035), and latitude (F = 1.71, *p* = 0.017) contributed significantly to the observed genetic variation. Partial RDA revealed that time significantly explained 4.5% of the total variance (F = 3.79, *p* = 0.001) while space significantly accounted for 2.0% (F = 1.64, *p* = 0.005) (**Table S11**). In *B. pomorum*, the combined effects of space and time explained 7.9% of the total genetic variance. Sequential tests indicated that longitude contributed significantly (F = 1.59, *p* = 0.009), while year and latitude were not significant. Partial RDA revealed that time explained 0.8% while space explained approximately 1.1% of the total variance, but both were not significant (**Table S11**).

#### 3.2.3 Linear mixed models (LMMs)

The correlation between the spatial and temporal variables was weak across species (Pearson correlation coefficient: *B. humilis* = 0.04, *B. sylvarum* = 0.09, *B. ruderatus* = 0.03, *B. veteranus* = 0.33, *B. pomorum* = 0.25), indicating low collinearity between predictors. Consequently, no multicollinearity correction was required, and both variables were considered independent in the mixed model formulation.

According to the LMMs (**Figure 6**, **Table S12**), the temporal distance had a significant positive effect on most species, indicating increased genetic divergence over time. The strongest positive effect of time on genetic distance was found in *B. ruderatus* (slope: 0.606, *p* < 2×10^-16^) where the high marginal R² value (41.9%), representing the variance explained only by fixed effects, indicates that temporal dynamics alone explained a substantial portion of the genetic variation observed. Temporal distance also had a significant positive effect in the genetic distance of *B. humilis* (slope: 0.228, *p* < 2×10^-16^), *B. sylvarum* (slope: 0.087, *p* = 0.041) and *B. veteranus* (slope: 0.151, *p* = 2.81×10^-04^). Geographical distance showed a significantly positive effect only on the genetic distance of *B. veteranus* (slope: 0.0531, *p* = 0.009) but had no significant effect on the genetic distance of the other species (**Figure 6**, **Table S12**). Finally, in *B. pomorum* no significant effects of temporal or spatial distances alone were observed on genetic distance, however a significant negative effect of their interaction was identified (slope: −0.0595, *p* = 0.026) (**Figure 6**, **Table S12**).

**Figure 6.**
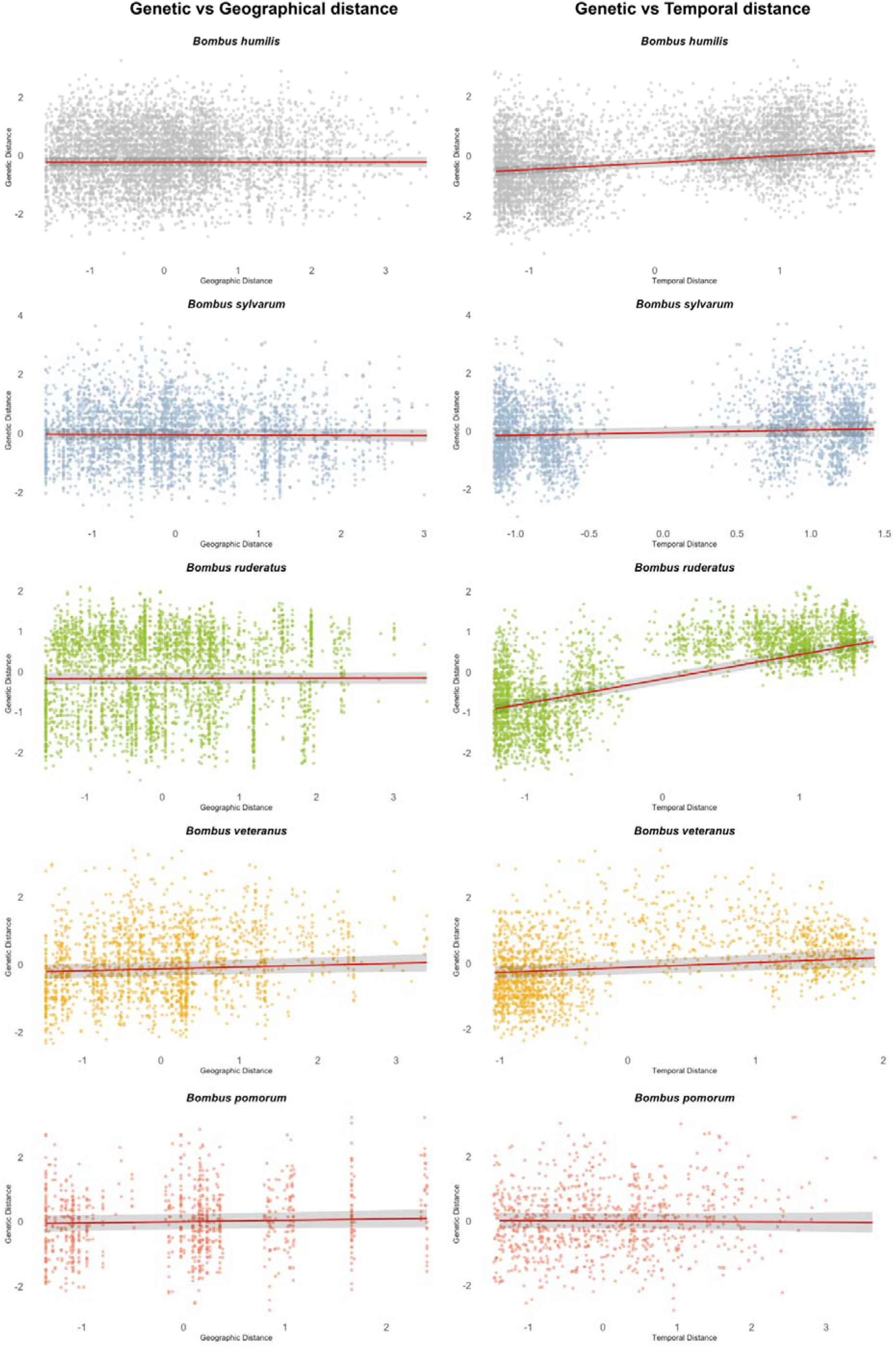
Spatial and temporal effect of genetic differentiation in each bumble bee species. For each species, the intercepts representing the linear correlation between genetic distance and spatial and temporal distance are displayed. A 95% of confidence interval has been also shown towards the intercept line.

In all species except *B. ruderatus*, low marginal R^2^ values (0.6%-5.4%) and high conditional R^2^ values (53.5%–84.1%), which account for both fixed and random effects, suggest that other factors rather than space and time might better explain the observed genetic variation (**Table S12**). Residuals followed a normal distribution with more than 95% of the values falling within the central quantile range (**Figure S6**).

## 4. Discussion

In the current study, the population genetics of five *Bombus* species undergoing different population dynamics over the past century was compared to investigate whether estimates of genetic diversity and population structure reflect occurrence data. Spatio-temporal population genetics revealed that genetic diversity indexes did not reflect occurrence patterns over time. However, shifts in allele frequencies were detected in three species (*B. ruderatus*, *B. humilis* and *B. veteranus*), which suggest recent colonization events in accordance with the occurrence data. Moreover, population fragmentation across space was detected in the cold-adapted *B. veteranus*, suggesting the presence of isolation by distance (IBD) and potential vulnerability in this species. In contrast, no IBD was observed in the extant warm-adapted species, suggesting stable gene flow among populations.

Reliance on museum collections constrained specimen availability and led to unequal sample sizes among the five *Bombus* species, particularly those with fluctuating or regionally extinct occurrence patterns. Relatively small sample sizes across an extended temporal scale likely contributed to the small effect sizes and low R² values observed in some models, reflecting inherent trade-offs in the study design. While denser sampling over a shorter time span might have increased statistical power to detect subtle or recent genetic changes, the broad temporal coverage used here allowed the assessment of longer-term trends. Nevertheless, the dataset provides a long-term perspective on genetic and demographic dynamics across species with contrasting population histories and highlights the importance of maintaining and expanding museum collections as critical resources for biodiversity monitoring, conservation, and retrospective genetic analyses.

### 4.1 Discrepancy between population occurrence and genetic diversity

Our findings revealed discrepancies between population occurrence and genetic diversity. For instance, despite demographic stability, *B. humilis* showed a significant decline in multilocus heterozygosity. In contrast, *B. ruderatus* maintained stable heterozygosity despite fluctuating occurrences, and *B. pomorum* exhibited slightly increased heterozygosity even as its population declined. Similar discrepancies between demography and genetic variation have been reported previously (Maebe et al., 2016; Rohde et al., 2024). Such patterns may reflect time lags between demographic events and genetic responses, during which the level of genetic diversity can persist despite the adverse conditions (Gargiulo et al., 2025). Additionally, the absence of specimens collected during population declines might have prevented the detection of potential genetic diversity losses occurring in these periods.

Nevertheless, despite substantial changes in population dynamics across those species, effect sizes associated with temporal changes in genetic diversity remained small, suggesting that the observed variation had limited biological relevance. Thus, genetic diversity indexes might not be the most adapted tools to monitor the conservation status of social bee populations. Small effect sizes and low R² values indicate that other parameters such as species-specific ecological traits and environmental factors other than time alone might influence both occurrence and genetic patterns (Romiguier et al., 2014). For instance, the disappearance of *B. pomorum* in Switzerland as well as in other European countries may be linked to its susceptibility to land-use changes, likely due to its late colony development cycle (Williams and Osborne, 2009). The species’ late male emergence and preference for thermophilic habitats, heavily impacted by human activities, probably contributed to its sharp decline. In Switzerland’s low-altitude warm regions, floral resources become scarce from late June onward, suggesting that mid-summer food limitation is likely a key driver of this species’ decline (https://species.infofauna.ch/groupe/1/conservation/149).

### 4.2 Temporal allelic shifts in Swiss bumble bee populations

Temporal allelic shifts were detected in *B. ruderatus*, *B. humilis* and *B. veteranus*. Even though such changes can result from several concurrent evolutionary processes (Stephan, 2016; Chen et al., 2019), the integration of genetic data with occurrence trends suggests that these allelic shifts are likely driven by gene flow from neighboring populations. The strongest evidence of such immigration was observed in *B. ruderatus*, which nearly vanished between 1960 and 1990, before reappearing from late 1990s/early 2000s with a markedly different allelic composition, consistent with recolonization. Similar processes may have acted in *B. humilis*, as this species showed a recent genetic shift likely attributed to immigration from northern populations primarily entering from the North-Eastern Swiss border, although less marked. However, while *B. ruderatus* near-extinction in northern Switzerland led to a complete allelic replacement, *B. humilis* remained stable over time, resulting in genetic admixture with immigrants. Subtler signals of allelic shift were also detected in *B. veteranus*. Historical occurrence data suggest that *B. veteranus* was once widespread across the Swiss Plateau but has recently shifted northward and colonized the Jura and the northern alpine Flanks, where there are no historical records of the species. Currently, the species is no longer found in its historical distribution range. Both genetic and occurrence data support the hypothesis of a recent colonization event, resulting in allelic shift, isolation by distance and change in the species’ distribution range. Additionally, this colonization may have contributed to the observed increased genetic diversity through admixture among different gene pools (Ingvarsson, 2001). However, limited contemporary specimens and the absence of recent material from the historical distribution range restrict the inference about genetic changes and demographic history. Applying genomic approaches would allow to detect selection signals and provide further insights into the factors that may be driving changes in bumble bee populations (Hart et al., 2022).

### 4.3 Effects of space and time on genetic differentiation in bumble bee populations

In contrast to time, space distance showed a weaker overall effect on genetic differentiation of warm-adapted *B. humilis*, *B. sylvarum* and *B. ruderatus*, indicating maintained gene flow and genetic connectivity over long distances (Cameron et al., 2011). However, spatial effects were weak but significant in regionally extinct and declining species. In *B. pomorum*, the negative interaction between geographical and temporal distance on genetic differentiation indicates increasing population homogenization before extinction. This pattern suggests that *B. pomorum* may have undergone a range contraction, during which originally separated populations were merged, resulting in genetic admixture and progressive homogenization that erased previous spatial genetic structure (Arenas et al., 2012). Over time, the loss of suitable habitats and the scarcity of floral resources may have driven this contraction, ultimately leading to the extinction of *B. pomorum* in the country. In contrast, the cold-adapted species (*B. veteranus*) showed a positive relationship between geographic and genetic distance, indicating isolation by distance (IBD). This pattern likely reflects recent shifts in the species’ distribution range, possibly driven by habitat fragmentation and loss (Jha, 2015; Müller and Praz, 2024) as well as climate change (Kerr et al., 2015; Martinet et al., 2021). Indeed, short-term changes in insect distributions have been linked to both climate change and regional land-use change, with warming trends strongly shaping long-term population dynamics in temperate regions (Neff et al., 2022). Based on the latest projections from Climate CH2025 (MeteoSwiss & ETH Zurich, 2025), Switzerland is warming faster than expected, with a temperature increase roughly twice the global average rate. Consistent with this pattern, many bumble bee species are undergoing population declines and range contractions (Nieto et al., 2014) and, as global temperatures rise, species might shift latitudinally and altitudinally to track suitable climatic conditions (Parmesan and Yohe, 2003; Kerr et al., 2015). Similarly, IBD patterns have been documented in other insects, including a threatened grasshopper species (Hoffmann et al., 2021) and in declining bumble bee species such as *B. pensylvanicus* (Lozier and Cameron, 2009) and *B. sylvarum* (Ellis et al., 2006).

Overall, small effect sizes and low marginal R^2^ values indicate that space and time only partially explained the observed genetic structure, suggesting that species-specific functional and ecological traits, such as phenology, foraging and habitat preferences, and demographic history likely play a substantial role in shaping species dynamics (Hofmann et al., 2019; Rasmont et al., 2021; Rohde et al., 2024).

## 5. Conclusions

Our results show that temporal changes in genetic diversity do not align with the magnitude of observed shifts in population dynamics in the five bumble bee species considered, suggesting that the genetic diversity indexes used here may not be adapted to assess the conservation status of social bees. Instead, the genetic data obtained here provides additional insights into species demographic history, revealing substantial migration processes that influenced population and genetic trajectories over time. Patterns of spatial genetic structure further highlight differences between extant warm- and cold-adapted species, with maintained gene flow in warm-adapted taxa contrasting with the detection of isolation by distance in the cold-adapted *B. veteranus*. The maintenance of gene flow despite drastic environmental changes in the region and period considered reflects the long-distance dispersal ability of bumble bees and suggests dynamic metapopulation processes. Thus, monitoring metapopulation dynamics in bumble bees might allow the detection of ongoing migration contributing to the persistence of populations. On the long term, ensuring connectivity among populations can be critical in facilitating gene flow and enabling demographic rescue processes, thereby supporting pollinator conservation at large spatial scales.

## Data availability statement

The list of specimens and relative information is included in the supplementary material file. Model specifications for R analyses are reported in the material and methods section. Full codes and genetic data will be made available upon acceptance.

## Acknowledgements

We would like to thank the Natural History Museums of Neuchatel, Bern, Lausanne and Fribourg and their curators Dr. Jessica Litman, Hannes Baur, Dr. Anne Freitag and Sophie Griens for granting us the access to the museum entomological collections. We also would like to thank Gaëlle Beureux and Giorgia Ferretti for helping with the lab work and contributing to the dataset. We would also thank the Research Institute of Organic Agriculture (FiBL) and Dr. Lucas-Barbosa Dani for providing specimens from their collection. We thank the Swiss data Centre Info Fauna for the access to the high-resolution dataset of the studied bumble bee species. We finally thank Marion Podolak for her support at the Natural History Museum of Lausanne and for providing the specimen pictures included in this publication.

## Authorship contribution statement

**Asia Piovesan:** Conceptualization, Data curation, Formal analysis, Investigation, Methodology, Software, Visualization, Writing-original draft, Writing-review and editing. **Christophe Praz:** Conceptualization, Data curation, Methodology, Validation, Writing-review and editing. **Bernhard Voelkl:** Formal analysis, Methodology, Software, Visualization, Writing-review and editing. **Sylvain Lanz:** Conceptualization, Data curation, Writing-original draft, Writing-review and editing. **Peter Neumann:** Funding acquisition, Resources, Project administration, Supervision, Writing-original draft, Writing-review and editing. **Alexis Beaurepaire:** Conceptualization, Investigation, Methodology, Project administration, Supervision, Validation, Writing-original draft, Writing-review and editing.

## Use of artificial intelligence

AI tools were used to assist with language editing during the manuscript preparation. All analyses, interpretations, and conclusions are the authors’ own.

