## Supplementary material for "Population genetics of bumble bee species with diverging population dynamics"

**SUPPLEMENTARY MATERIALS**

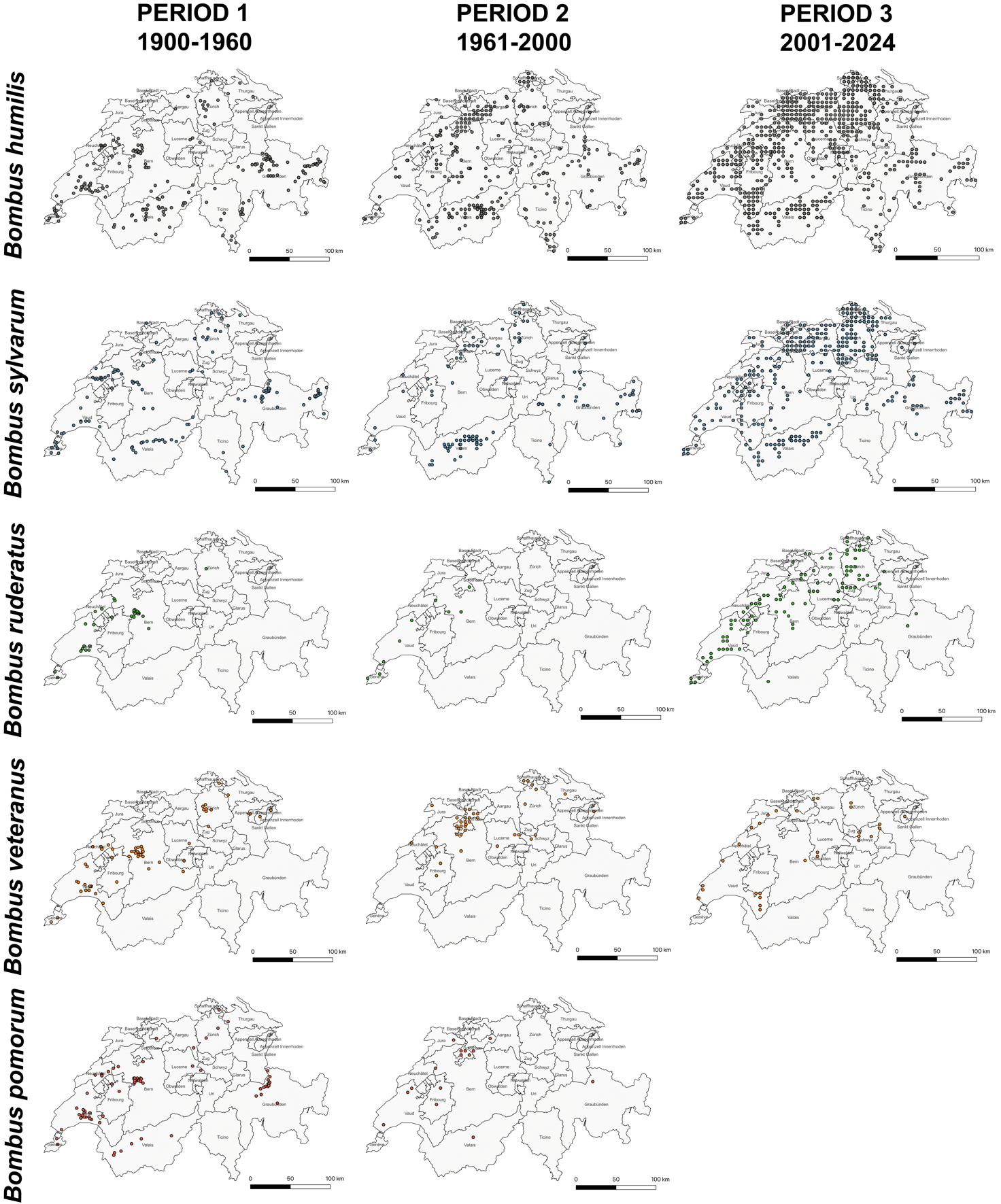

**Figure S1**. **Occurrence maps of the five species in Switzerland**. Georeferenced data of specimens collected in Switzerland for each species from 1900 to 2024. For each species three time periods were defined (PERIOD 1: 1900-1960, PERIOD 2: 1961-2000 and PERIOD 3: 2001-2024). The location of specimens collected in the corresponding time periods are represented in the map.

**Table S1. List of specimens of *B. humilis*, *B. sylvarum*, *B. ruderatus*, *B. veteranus* and *B. pomorum*.** In the table are reported the specimens genotyped for each species with the correspondent GBIF number, the year and sampling locality with GPS coordinates (LATITUDE and LONGITUDE). The museum where the specimen is housed is also provided. The decade to which each specimen has been assigned has been reported through the cluster number assignment. In *B. veteranus* a * was used to indicate the specimen excluded from the analyses after sibship inference.

| **SPECIES** | **GBIF** | **YEAR** | **CLUSTER** | **SAMPLING LOCALITY** | **LATITUDE** | **LONGITUDE** | **MUSEUM** |
| --- | --- | --- | --- | --- | --- | --- | --- |
| *B. humilis* | GBIFCH00412278 | 1929 | 1 | b. Treiten | 47.00920 | 7.15590 | NMBE |
| *B. humilis* | GBIFCH00412265 | 1929 | 1 | Vinelz | 47.03608 | 7.11628 | NMBE |
| *B. humilis* | GBIFCH00412261 | 1930 | 2 | Aventicum | 46.88292 | 7.03849 | NMBE |
| *B. humilis* | GBIFCH00412362 | 1930 | 2 | Selhofen | 46.91960 | 7.47145 | NMBE |
| *B. humilis* | GBIFCH00232772 | 1932 | 2 | Vaud, Buchillon | 46.47433 | 6.42952 | MZL |
| *B. humilis* | GBIFCH00232847 | 1932 | 2 | Epalinges | 46.55722 | 6.67576 | MZL |
| *B. humilis* | GBIFCH00232805 | 1933 | 2 | Genève, Cologny | 46.21987 | 6.17503 | MZL |
| *B. humilis* | GBIFCH00232913 | 1933 | 2 | Nyon | 46.38248 | 6.23624 | MZL |
| *B. humilis* | GBIFCH00233285 | 1933 | 2 | Genève, Chambésy | 46.24656 | 6.14847 | MZL |
| *B. humilis* | GBIFCH00232787 | 1933 | 2 | Neuchâtel, Auvernier | 46.97220 | 6.88012 | MZL |
| *B. humilis* | GBIFCH00412122 | 1934 | 2 | Zollikofen | 47.00055 | 7.45835 | NMBE |
| *B. humilis* | GBIFCH00412139 | 1934 | 2 | Muri | 46.92859 | 7.48459 | NMBE |
| *B. humilis* | GIBIFCH00118187 | 1935 | 2 | Bois de Veyrier | 46.16606 | 6.18921 | MHNG |
| *B. humilis* |  | 1935 | 2 | Mont Vully | 46.96405 | 7.09045 | MHNF |
| *B. humilis* |  | 1935 | 2 | Yverdon | 46.78192 | 6.64639 | NMSH |
| *B. humilis* | GBIFCH00232815 | 1936 | 2 | Lausanne, Bois de Belmont | 46.52124 | 6.67626 | MZL |
| *B. humilis* | GBIFCH00233243 | 1936 | 2 | Vaud, St. Sulpice | 46.51129 | 7.67626 | MZL |
| *B. humilis* | GBIFCH00233265 | 1938 | 2 | Vaud, Cully | 46.48560 | 6.72886 | MZL |
| *B. humilis* | GBIFCH00232966 | 1939 | 2 | Lausanne, Bois de Belmont | 46.52124 | 6.67626 | MZL |
| *B. humilis* | GBIFCH00233064 | 1940 | 3 | Neuchâtel, Cortaillod | 46.94502 | 6.84100 | MZL |
| *B. humilis* | GBIFCH00233154 | 1941 | 3 | Lausanne, Bois de Belmont | 46.52124 | 6.67627 | MZL |
| *B. humilis* | GBIFCH00233328 | 1941 | 3 | VD, Yvonand | 46.80058 | 6.75091 | MZL |
| *B. humilis* | GBIFCH00233282 | 1942 | 3 | Vaud, Puidoux | 46.50390 | 6.78074 | MZL |
| *B. humilis* | GBIFCH00233272 | 1942 | 3 | Neuchâtel, Auvernier | 46.97220 | 6.88012 | MZL |
| *B. humilis* | GBIFCH00232896 | 1943 | 3 | Jorat | 46.58437 | 6.70147 | MZL |
| *B. humilis* | GBIFCH00232804 | 1943 | 3 | Cossonay | 46.60992 | 6.50531 | MZL |
| *B. humilis* | GBIFCH00232906 | 1943 | 3 | Vaud, Romanel | 46.56566 | 6.59738 | MZL |
| *B. humilis* | GBIFCH00232791 | 1944 | 3 | Caux, Naye | 46.43273 | 6.93773 | MZL |
| *B. humilis* | GBIFCH00232922 | 1944 | 3 | Neuchâtel, Auvernier | 46.97220 | 6.88012 | MZL |
| *B. humilis* | GBIFCH00233069 | 1945 | 3 | Chalet-à-Gobet | 46.56630 | 6.68868 | MZL |
| *B. humilis* | GBIFCH00232799 | 1945 | 3 | Vaud, St. Sulpice | 46.51129 | 6.54611 | MZL |
| *B. humilis* | GBIFCH00233129 | 1945 | 3 | Neuchâtel, Auvernier | 46.97220 | 6.88012 | MZL |
| *B. humilis* | GBIFCH00233073 | 1946 | 3 | Cologny | 46.21987 | 6.17503 | MZL |
| *B. humilis* | GBIFCH00232769 | 1946 | 3 | Vaud, St. Sulpice | 46.51129 | 6.54611 | MZL |
| *B. humilis* | GBIFCH00232826 | 1946 | 3 | Neuchâtel, Auvernier | 46.97220 | 6.88012 | MZL |
| *B. humilis* | GBIFCH00232984 | 1947 | 3 | Lausanne, Bois de Belmont | 46.52124 | 6.67627 | MZL |
| *B. humilis* | GBIFCH00233252 | 1947 | 3 | Cologny | 46.21987 | 6.17503 | MZL |
| *B. humilis* | GBIFCH00232773 | 1947 | 3 | Neuchâtel, Auvernier | 46.97220 | 6.88012 | MZL |
| *B. humilis* | GBIFCH00232807 | 1947 | 3 | Neuchâtel, Auvernier | 46.97220 | 6.88012 | MZL |
| *B. humilis* | GBIFCH00328099 | 1948 | 3 | Bern | 46.94659 | 7.44520 | NMBE |
| *B. humilis* | GBIFCH00233050 | 1948 | 3 | Neuchâtel, Corcelles | 46.98120 | 6.88003 | MZL |
| *B. humilis* | GBIFCH00326198 | 1948 | 3 | Neuchâtel | 46.99943 | 6.93243 | NMLU |
| *B. humilis* | GBIFCH00232921 | 1949 | 3 | Vaud, Vaux s/Morges | 46.53763 | 6.46745 | MZL |
| *B. humilis* | GBIFCH00412196 | 1949 | 3 | Bern, Elfenau BE | 46.92859 | 7.47146 | NMBE |
| *B. humilis* | GBIFCH00328137 | 1950 | 4 | Bern, Bethlehem | 46.94658 | 7.39266 | NMBE |
| *B. humilis* | GBIFCH00328119 | 1951 | 4 | Belpmoos BE | 46.91059 | 7.49770 | NMBE |
| *B. humilis* | GBIFCH00328175 | 1952 | 4 | Rehhag BE | 46.93757 | 7.37953 | NMBE |
| *B. humilis* | GBIFCH00232800 | 1955 | 4 | Suisse, Genève, Cartigny | 46.17305 | 6.02071 | MZL |
| *B. humilis* | GBIFCH00412247 | 1984 | 5 | BREMGARTEN | 46.98257 | 7.44520 | NMBE |
| *B. humilis* | GBIFCH00328251 | 1986 | 5 | Onnens VD | 46.83615 | 6.68491 | NMBE |
| *B. humilis* | GBIFCH00328255 | 1989 | 5 | Vauffelin Füglisthal (BE) | 47.18936 | 7.30008 | NMBE |
| *B. humilis* | GBIFCH00412235 | 1990 | 6 | BELP | 46.92859 | 7.47146 | NMBE |
| *B. humilis* | GBIFCH00412204 | 1991 | 6 | BELP | 46.92859 | 7.47146 | NMBE |
| *B. humilis* | GBIFCH00412174 | 1993 | 6 | OSCHWALD | 47.14416 | 7.70889 | NMBE |
| *B. humilis* | GBIFCH00328273 | 1997 | 6 | BELP | 46.92859 | 7.47146 | NMBE |
| *B. humilis* | GBIFCH00434023 | 1998 | 6 | VD, Eclépens | 46.65531 | 6.55679 | NMBE |
| *B. humilis* | GBIFCH00434033 | 1998 | 6 | NE, Noiraigue | 46.97149 | 6.74871 | NMBE |
| *B. humilis* | GBIFCH00328282 | 1998 | 6 | LIEBEFELD | 46.92859 | 7.40581 | NMBE |
| *B. humilis* | GBIFCH00328246 | 1999 | 6 | BELP | 46.92859 | 7.47146 | NMBE |
| *B. humilis* | GBIFCH00328263 | 1999 | 6 | LIEBEFELD | 46.92859 | 7.40581 | NMBE |
| *B. humilis* | GBIFCH00328307 | 2000 | 7 | BELP | 46.92859 | 7.47146 | NMBE |
| *B. humilis* | GBIFCH00412331 | 2000 | 7 | VD, Sergey | 46.74473 | 6.48989 | NMBE |
| *B. humilis* | GBIFCH00328335 | 2002 | 7 | VD, Romainmôtier | 46.69920 | 6.42531 | NMBE |
| *B. humilis* | GBIFCH00412161 | 2002 | 7 | VD, Champagne | 46.83615 | 6.68491 | NMBE |
| *B. humilis* | GBIFCH00328300 | 2003 | 7 | INS | 46.97319 | 7.14295 | NMBE |
| *B. humilis* | GBIFCH00412167 | 2003 | 7 | GE, Chancy | 46.15456 | 5.98235 | NMBE |
| *B. humilis* | GBIFCH00412189 | 2003 | 7 | Pèry | 47.19832 | 7.27367 | NMBE |
| *B. humilis* | GBIFCH00412257 | 2003 | 7 | LIEBEFELD | 46.92859 | 7.40581 | NMBE |
| *B. humilis* | GBIFCH00328295 | 2004 | 7 | BE, Rüeschegg | 46.79366 | 7.40589 | NMBE |
| *B. humilis* | GBIFCH00412178 | 2004 | 7 | WIEDLISBACH | 47.24330 | 7.60373 | NMBE |
| *B. humilis* | GBIFCH00412181 | 2004 | 7 | KRÄILIGEN | 47.15343 | 7.53752 | NMBE |
| *B. humilis* | GBIFCH00412253 | 2004 | 7 | LIEBEFELD | 46.92859 | 7.40581 | NMBE |
| *B. humilis* | GBIFCH00328324 | 2005 | 7 | GRANGES | 46.79196 | 6.81651 | NMBE |
| *B. humilis* | GBIFCH00328314 | 2006 | 7 | GE, Russin | 46.19953 | 5.98115 | NMBE |
| *B. humilis* | GBIFCH0063619 | 2013 | 8 | Orvin | 47.15324 | 7.20788 | MHNN |
| *B. humilis* | GBIFCH00118866 | 2013 | 8 | Moutier, Les Golats | 47.28838 | 7.40558 | MHNN |
| *B. humilis* | GBIFCH00118906 | 2013 | 8 | Neuchâtel, Jardin Botanique | 47.01748 | 6.94541 | MHNN |
| *B. humilis* | GBIFCH00093522 | 2014 | 8 | Champvent, St.-Christophe | 46.79023 | 6.55458 | MHNN |
| *B. humilis* | GBFCH0499686 | 2014 | 8 | Cossonay-Ville, Le Signal | 46.60420 | 6.50959 | MHNN |
| *B. humilis* | GBIFCH00100514 | 2014 | 8 | Moulin-de-Vert | 46.20035 | 6.04591 | MHNN |
| *B. humilis* | GBFCH049471 | 2014 | 8 | Heiligenschwendi | 46.75181 | 7.67912 | MHNN |
| *B. humilis* | GBIFCH049501 | 2014 | 8 | Arnex-sur-Orbe, Les Vaux | 46.69976 | 6.49068 | MHNN |
| *B. humilis* | GBIFCH0063265 | 2014 | 8 | Le Landeron | 47.06295 | 7.07663 | MHNN |
| *B. humilis* | GBIFCH00410403 | 2015 | 8 | La Mauguettaz, Les Vursys | 46.79159 | 6.75102 | MZL |
| *B. humilis* | GBIFCH00121254 | 2016 | 8 | Val-de-Travers, Couvet | 46.92565 | 6.61800 | MHNN |
| *B. humilis* | GBIFCH0072190 | 2016 | 8 | La Mauguettaz | 46.79159 | 6.75102 | MHNN |
| *B. humilis* |  | 2016 | 8 | Wynau | 47.26050 | 7.80098 | FIBL |
| *B. humilis* |  | 2016 | 8 | Toffen | 46.84878 | 7.49975 | FIBL |
| *B. humilis* | GBIFCH00118783 | 2016 | 8 | Grandval | 47.28838 | 7.40558 | MHNN |
| *B. humilis* | GBIFCH00499039 | 2016 | 8 | Lausanne, Beaulieu | 46.52087 | 6.62414 | MZL |
| *B. humilis* | GBIFCH00123257 | 2016 | 8 | Noville, Les Grangettes | 46.38751 | 6.88615 | MHNN |
| *B. humilis* | GBIFCH00127919 | 2017 | 8 | Farnern | 47.28827 | 7.60387 | MHNN |
| *B. humilis* | GBIFCH00119491 | 2017 | 8 | Gampelen, Rothaus | 47.01775 | 7.01117 | MHNN |
| *B. humilis* | GBIFCH00132167 | 2017 | 8 | Monts-de-Corsier | 46.52245 | 6.88477 | MHNN |
| *B. humilis* | GBIFCH00119985 | 2018 | 8 | Eclépens, Mormont | 46.65531 | 6.55679 | MHNN |
| *B. humilis* | GBIFCH00117579 | 2018 | 8 | Gondiswil | 47.15267 | 7.86716 | MHNN |
| *B. humilis* | GBFCH00093504 | 2019 | 8 | Orny | 46.66308 | 6.55432 | MHNN |
| *B. humilis* | GBFCH00093534 | 2019 | 8 | Onnens | 46.85107 | 6.69092 | MHNN |
| *B. humilis* | GBIFCH00452967 | 2019 | 8 | Neuchâtel | 47.01748 | 6.94541 | MHNF |
| *B. humilis* | GBIFCH00136499 | 2019 | 8 | Les Ponts-de-Martel | 46.97107 | 6.68301 | MHNN |
| *B. humilis* | GBIFCH00136650 | 2019 | 8 | Chancy | 46.15456 | 5.98235 | MHNN |
| *B. humilis* | GBIFCH00129309 | 2020 | 9 | Därstetten, Gruebi | 46.65869 | 7.53662 | MHNN |
| *B. humilis* | GBIFCH00991454 | 2020 | 9 | Leysin, N La Roulaz | 46.34310 | 7.01650 | MZL |
| *B. humilis* | GBIFCH049324 | 2021 | 9 | La Sarraz | 46.65479 | 6.49147 | MHNN |
| **SPECIES** | **GBIF** | **YEAR** | **CLUSTER** | **SAMPLING LOCALITY** | **LATITUDE** | **LONGITUDE** | **MUSEUM** |
| *B. sylvarum* | GBIFCH00427057 | 1929 | 1 | Wohlen, Lörmoos | 46.98215 | 7.34786 | MZL |
| *B. sylvarum* | GBIFCH00427075 | 1929 | 1 | Meiringen | 46.72848 | 8.18278 | MZL |
| *B. sylvarum* | GBIFCH00236986 | 1930 | 2 | Genève, La Plaine | 46.18171 | 5.99458 | MZL |
| *B. sylvarum* | GBIFCH00236977 | 1931 | 2 | Vaud, Buchillon | 46.47433 | 6.42952 | MZL |
| *B. sylvarum* | GBIFCH00271176 | 1931 | 2 | Hte Savoie, Pied du Salève | 46.14791 | 6.17670 | MZL |
| *B. sylvarum* | GBIFCH00236865 | 1932 | 2 | Genève, Cologny | 46.21987 | 6.17503 | MZL |
| *B. sylvarum* | GBIFCH00236928 | 1932 | 2 | Genève, Cologny | 46.21987 | 6.17503 | MZL |
| *B. sylvarum* | GBIFCH00236942 | 1932 | 2 | Vaud, Promenthoux | 46.39188 | 6.27504 | MZL |
| *B. sylvarum* | GBIFCH00271181 | 1932 | 2 | Hte Savoie, Ballaison | 46.30245 | 6.32888 | MZL |
| *B. sylvarum* | GBIFCH00271198 | 1932 | 2 | Hte Savoie, Sciez | 46.32993 | 6.38026 | MZL |
| *B. sylvarum* | GBIFCH00271204 | 1932 | 2 | Hte Savoie, Pied du Salève | 46.14791 | 6.17670 | MZL |
| *B. sylvarum* | GBIFCH00271143 | 1932 | 2 | Hte Savoie, Ballaison | 46.30245 | 6.32888 | MZL |
| *B. sylvarum* | GBIFCH00271155 | 1932 | 2 | Hte Savoie, Salève | 46.13906 | 6.18984 | MZL |
| *B. sylvarum* | GBIFCH00237536 | 1932 | 2 | Genève, Cologny | 46.21987 | 6.17503 | MZL |
| *B. sylvarum* | GBIFCH00232748 | 1933 | 2 | Genève, Chambésy | 46.24656 | 6.14847 | MZL |
| *B. sylvarum* | GBIFCH00232751 | 1933 | 2 | Neuchâtel, Auvernier | 46.97220 | 6.88012 | MZL |
| *B. sylvarum* | GBIFCH00232749 | 1933 | 2 | Genève, Cologny | 46.21987 | 6.17503 | MZL |
| *B. sylvarum* | GBIFCH00232754 | 1933 | 2 | Genève, Chambésy | 46.24656 | 6.14847 | MZL |
| *B. sylvarum* | GBIFCH00232760 | 1933 | 2 | Neuchâtel, Auvernier | 46.97220 | 6.88012 | MZL |
| *B. sylvarum* | GBIFCH00236872 | 1933 | 2 | Neuchâtel, Auvernier | 46.97220 | 6.88012 | MZL |
| *B. sylvarum* | GBIFCH00236859 | 1933 | 2 | Neuchâtel, Auvernier | 46.97220 | 6.88012 | MZL |
| *B. sylvarum* | GBIFCH00236867 | 1933 | 2 | Neuchâtel, Auvernier | 46.97220 | 6.88012 | MZL |
| *B. sylvarum* | GBIFCH00236966 | 1933 | 2 | Vaud, Chéserex | 46.39975 | 6.17083 | MZL |
| *B. sylvarum* | GBIFCH00236948 | 1933 | 2 | Nyon | 46.38248 | 6.23624 | MZL |
| *B. sylvarum* | GBIFCH00236950 | 1933 | 2 | Venoge, St-Sulpice VD | 46.51129 | 6.54611 | MZL |
| *B. sylvarum* | GBIFCH00271199 | 1933 | 2 | Hte Savoie, Salève | 46.14791 | 6.17670 | MZL |
| *B. sylvarum* | GBIFCH00237512 | 1933 | 2 | Vaud, Giez | 46.81388 | 6.61007 | MZL |
| *B. sylvarum* | GBIFCH00237543 | 1933 | 2 | Genève, Bernex | 46.18021 | 6.06999 | MZL |
| *B. sylvarum* | GBIFCH00431489 | 1934 | 2 | St-Blaise | 47.01764 | 6.98487 | MZL |
| *B. sylvarum* | GBIFCH00427013 | 1934 | 2 | St-Blaise | 47.01570 | 6.98746 | MZL |
| *B. sylvarum* | GBIFCH00427023 | 1934 | 2 | Bern | 46.95456 | 7.42069 | MZL |
| *B. sylvarum* | GBIFCH00237517 | 1934 | 2 | Vaud, St. Sulpice | 46.51238 | 6.56026 | MZL |
| *B. sylvarum* | GBIFCH00236868 | 1935 | 2 | Vaud, St. Sulpice | 46.51129 | 6.54611 | MZL |
| *B. sylvarum* | GBIFCH00271182 | 1935 | 2 | Hte Savoie, Pied du Salève | 46.14791 | 6.17670 | MZL |
| *B. sylvarum* | GBIFCH00237539 | 1935 | 2 | Vaud, St. Sulpice | 46.51238 | 6.56026 | MZL |
| *B. sylvarum* | GBIFCH00271144 | 1935 | 2 | Hte Savoie, Pied du Salève | 46.51238 | 6.56026 | MZL |
| *B. sylvarum* | GBIFCH00271156 | 1935 | 2 | Hte Savoie, Pied du Salève | 46.51238 | 6.56026 | MZL |
| *B. sylvarum* | GBIFCH00236862 | 1936 | 2 | Vaud, St. Sulpice | 46.51129 | 6.54611 | MZL |
| *B. sylvarum* | GBIFCH00236869 | 1936 | 2 | Vaud, St. Sulpice | 46.51129 | 6.54611 | MZL |
| *B. sylvarum* | GBIFCH00236875 | 1936 | 2 | Vaud, St. Sulpice | 46.51129 | 6.54611 | MZL |
| *B. sylvarum* | GBIFCH00237532 | 1936 | 2 | Vaud, St. Sulpice | 46.51238 | 6.56026 | MZL |
| *B. sylvarum* | GBIFCH00237530 | 1936 | 2 | Vaud, St. Sulpice | 46.51238 | 6.56026 | MZL |
| *B. sylvarum* | GBIFCH00237525 | 1936 | 2 | Vaud, St. Sulpice | 46.51238 | 6.56026 | MZL |
| *B. sylvarum* | GBIFCH00237523 | 1936 | 2 | Vaud, St. Sulpice | 46.51238 | 6.56026 | NMBE |
| *B. sylvarum* | GBIFCH00237542 | 1939 | 2 | Vaud, La Chaux (Cossonay) | 46.62373 | 6.46812 | NMBE |
| *B. sylvarum* | GBIFCH00237562 | 1939 | 2 | Neuchâtel, Auvernier | 46.97220 | 6.88012 | NMBE |
| *B. sylvarum* | GBIFCH00237534 | 1940 | 3 | Vaud, Noville | 46.39379 | 6.89089 | NMBE |
| *B. sylvarum* | GBIFCH00427004 | 1940 | 3 | Bern, Tierpark | 46.93460 | 7.45110 | NMBE |
| *B. sylvarum* | GBIFCH00426966 | 1940 | 3 | Bern | 46.95456 | 7.42069 | NMBE |
| *B. sylvarum* | GBIFCH00427019 | 1940 | 3 | Bern, Tierpark | 46.93460 | 7.45110 | NMBE |
| *B. sylvarum* | GBIFCH00431438 | 1999 | 4 | BELP | 46.92859 | 7.47146 | NMBE |
| *B. sylvarum* | GBIFCH00431476 | 2005 | 5 | MISERY | 46.83807 | 7.07815 | NMBE |
| *B. sylvarum* | GBIFCH00118896 | 2013 | 6 | La Sarraz, Le Signal | 46.65479 | 6.49147 | MHNN |
| *B. sylvarum* | GBIFCH049198 | 2014 | 6 | Bonvillars | 46.83615 | 6.68491 | MHNN |
| *B. sylvarum* | GBIFCH00093497 | 2014 | 6 | Lucens, Rive Ouest (L) | 46.70235 | 6.88292 | MHNN |
| *B. sylvarum* | GBIFCH00092977 | 2014 | 6 | Orny | 46.65479 | 6.49147 | MHNN |
| *B. sylvarum* | GIBFCH00093692 | 2014 | 6 | Longirod, Bugnonet | 46.50157 | 6.26603 | MHNN |
| *B. sylvarum* | GBIFCH00093475 | 2014 | 6 | Bonvillars | 46.83615 | 6.68491 | MHNN |
| *B. sylvarum* | GBIFCH00101494 | 2014 | 6 | Moulin-de-Vert | 46.20035 | 6.04591 | MHNN |
| *B. sylvarum* | GBIFCH00118796 | 2016 | 6 | Röthenbach im Emmental | 46.83808 | 7.79912 | MHNN |
| *B. sylvarum* | GBIFCH00123264 | 2016 | 6 | Noville, Les Grangettes | 46.38751 | 6.88615 | MHNN |
| *B. sylvarum* | GBIFCH00127908 | 2017 | 6 | Farnern | 47.28827 | 7.60387 | MHNN |
| *B. sylvarum* | GBIFCH00121614 | 2017 | 6 | Chardonne | 46.47747 | 6.88523 | MHNN |
| *B. sylvarum* | GBIFCH00132176 | 2017 | 6 | Monts-de-Corsier | 46.52245 | 6.88477 | MHNN |
| *B. sylvarum* | GBIFCH00135596 | 2018 | 6 | Ferreyres | 46.65479 | 6.49147 | MHNN |
| *B. sylvarum* | GBIFCH00127958 | 2018 | 6 | Heimberg | 46.79355 | 7.60236 | MHNN |
| *B. sylvarum* | GBIFCH00139330 | 2018 | 6 | St-Blaise | 47.01775 | 7.01117 | MHNN |
| *B. sylvarum* | GIBFCH00092893 | 2019 | 6 | Trélex | 46.43100 | 6.18445 | MHNN |
| *B. sylvarum* | GBIFCH00093377 | 2019 | 6 | Eclépens | 46.65531 | 6.55679 | MHNN |
| *B. sylvarum* | GBIFCH00093265 | 2019 | 6 | Trélex | 46.42674 | 6.17020 | MHNN |
| *B. sylvarum* | GBIFCH00129528 | 2019 | 6 | Château-d¿Oex | 46.47825 | 7.08055 | MHNN |
| *B. sylvarum* | GBIFCH00123496 | 2019 | 6 | Bourg-en-Lavaux, Les Auges | 46.52173 | 6.75446 | MHNN |
| *B. sylvarum* | GBIFCH00123497 | 2019 | 6 | Bourg-en-Lavaux, Les Auges | 46.52173 | 6.75446 | MHNN |
| *B. sylvarum* | GBIFCH00129430 | 2020 | 7 | Därstetten, Nidfluh | 46.65869 | 7.53662 | MHNN |
| *B. sylvarum* | GBIFCH00093084 | 2021 | 7 | Yens | 46.51930 | 6.42868 | MHNN |
| *B. sylvarum* | GBIFCH049519 | 2021 | 7 | VInzel | 46.42812 | 6.30028 | MHNN |
| *B. sylvarum* | GBIFCH00093004 | 2021 | 7 | Bursins | 46.42812 | 6.30028 | MHNN |
| **SPECIES** | **GBIF** | **YEAR** | **CLUSTER** | **SAMPLING LOCALITY** | **LATITUDE** | **LONGITUDE** | **MUSEUM** |
| *B. ruderatus* | GBIFCH00426862 | 1930 | 1 | Aventicum | 46.88292 | 7.03849 | NMBE |
| *B. ruderatus* | NMB-HYMEN0000168 | 1930 | 1 | Bern Dählhölzli | 46.93759 | 7.45833 | Bâle |
| *B. ruderatus* | GBIFCH00192755 | 1933 | 1 | Neuchâtel, Auvernier | 46.9722 | 6.88012 | MZL |
| *B. ruderatus* | GBIFCH00192756 | 1933 | 1 | Neuchâtel, Auvernier | 46.9722 | 6.88012 | MZL |
| *B. ruderatus* | GBIFCH00192770 | 1933 | 1 | Neuchâtel, Auvernier | 46.9722 | 6.88012 | MZL |
| *B. ruderatus* | GBIFCH00192829 | 1933 | 1 | Neuchâtel, Auvernier | 46.9722 | 6.88012 | MZL |
| *B. ruderatus* | GBIFCH00193123 | 1933 | 1 | Neuchâtel, Auvernier | 46.9722 | 6.88012 | MZL |
| *B. ruderatus* | GBIFCH00193127 | 1933 | 1 | Neuchâtel, Auvernier | 46.9722 | 6.88012 | MZL |
| *B. ruderatus* | GBIFCH00193264 | 1933 | 1 | Neuchâtel, Auvernier | 46.9722 | 6.88012 | MZL |
| *B. ruderatus* | GBIFCH00192783 | 1933 | 1 | Genève, Cologny | 46.21987 | 6.17503 | MZL |
| *B. ruderatus* | GBIFCH00192816 | 1933 | 1 | Genève, Cologny | 46.21987 | 6.17503 | MZL |
| *B. ruderatus* | GBIFCH00193167 | 1933 | 1 | Lausanne | 46.52087 | 6.62414 | MZL |
| *B. ruderatus* | GBIFCH00192824 | 1933 | 1 | Genève, Cologny | 46.21987 | 6.17503 | MZL |
| *B. ruderatus* | GBIFCH00193117 | 1933 | 1 | Vaud, Giez | 46.80871 | 6.61978 | MZL |
| *B. ruderatus* | GBIFCH00193289 | 1933 | 1 | Vaud, Giez | 46.80871 | 6.61978 | MZL |
| *B. ruderatus* | GBIFCH00426876 | 1934 | 1 | Bern | 46.94659 | 7.4452 | NMBE |
| *B. ruderatus* | GBIFCH00193181 | 1935 | 1 | Vaud, St. Sulpice | 46.51129 | 6.54611 | MZL |
| *B. ruderatus* | GBIFCH00192831 | 1935 | 1 | Neuchâtel, Auvernier | 46.9722 | 6.88012 | MZL |
| *B. ruderatus* | GBIFCH00193134 | 1936 | 1 | Lausanne, Bois de Belmont | 46.52124 | 6.67627 | MZL |
| *B. ruderatus* | GBIFCH00193177 | 1940 | 2 | Vaud, Yvonand | 46.80058 | 6.75091 | MZL |
| *B. ruderatus* | GBIFCH00192760 | 1941 | 2 | VD, Yvonand | 46.80058 | 6.75091 | MZL |
| *B. ruderatus* | GBIFCH00192833 | 1941 | 2 | VD, Yvonand | 46.80058 | 6.75091 | MZL |
| *B. ruderatus* | GBIFCH00192836 | 1942 | 2 | Neuchâtel, Auvernier | 46.9722 | 6.88012 | MZL |
| *B. ruderatus* | GBIFCH00192815 | 1942 | 2 | Neuchâtel, Auvernier | 46.9722 | 6.88012 | MZL |
| *B. ruderatus* | GBIFCH00193156 | 1943 | 2 | St. Sulpice VD | 46.51129 | 6.54611 | MZL |
| *B. ruderatus* | GBIFCH00192888 | 1944 | 2 | Neuchâtel, Auvernier | 46.9722 | 6.88012 | MZL |
| *B. ruderatus* | GBIFCH00426888 | 1944 | 2 | Liebefeld | 46.92859 | 7.41894 | NMBE |
| *B. ruderatus* | GBIFCH00192801 | 1944 | 2 | St. Sulpice VD | 46.51129 | 6.54611 | MZL |
| *B. ruderatus* | GBIFCH00192788 | 1945 | 2 | Lausanne, Romanel | 46.56566 | 6.59738 | MZL |
| *B. ruderatus* | GBIFCH00192806 | 1945 | 2 | Suisse, Vaud, St. Sulpice | 46.51129 | 6.54611 | MZL |
| *B. ruderatus* | GBIFCH00192808 | 1945 | 2 | Cologny | 46.21987 | 6.17503 | MZL |
| *B. ruderatus* | GBIFCH00192823 | 1945 | 2 | Lausanne, Romanel | 46.56566 | 6.59738 | MZL |
| *B. ruderatus* | GBIFCH00192886 | 1945 | 2 | Cologny | 46.21987 | 6.17503 | MZL |
| *B. ruderatus* | GBIFCH00193148 | 1945 | 2 | Cologny | 46.21987 | 6.17503 | MZL |
| *B. ruderatus* | GBIFCH00193162 | 1945 | 2 | Lausanne, Romanel | 46.56566 | 6.59738 | MZL |
| *B. ruderatus* | GBIFCH00192769 | 1946 | 2 | Genève, Cologny | 46.21987 | 6.17503 | MZL |
| *B. ruderatus* | GBIFCH00192837 | 1946 | 2 | Genève, Cologny | 46.21987 | 6.17503 | MZL |
| *B. ruderatus* | GBIFCH00426877 | 1948 | 2 | Bern, Bümpliz | 46.93758 | 7.39267 | NMBE |
| *B. ruderatus* | GBIFCH00192752 | 1948 | 2 | Auvernier | 46.9722 | 6.88012 | MZL |
| *B. ruderatus* | GBIFCH00426850 | 1951 | 3 | Sportplatz Wander BE | 46.93759 | 7.41893 | NMBE |
| *B. ruderatus* | GBIFCH00426886 | 1951 | 3 | Hunziken BE | 46.89257 | 7.53705 | NMBE |
| *B. ruderatus* | GBIFCH00426895 | 1952 | 3 | Landorf BE | 46.91959 | 7.40581 | NMBE |
| *B. ruderatus* | GBIFCH00426885 | 1952 | 3 | Bremgartenwald BE | 46.96457 | 7.40578 | NMBE |
| *B. ruderatus* | GBIFCH00426899 | 1953 | 3 | Bethleh., b. Messerligr. BE | 46.95557 | 7.39265 | NMBE |
| *B. ruderatus* | GBIFCH00426867 | 1953 | 3 | Messerligrube BE | 46.95557 | 7.39265 | NMBE |
| *B. ruderatus* | GBIFCH00426883 | 1990 | 4 | BELP | 46.92859 | 7.47146 | NMBE |
| *B. ruderatus* | GBIFCH00426890 | 1991 | 4 | BELP | 46.92859 | 7.47146 | NMBE |
| *B. ruderatus* | GBIFCH00426896 | 1995 | 4 | BELP | 46.92859 | 7.47146 | NMBE |
| *B. ruderatus* | GBIFCH00426833 | 1997 | 4 | BELP | 46.92859 | 7.47146 | NMBE |
| *B. ruderatus* | GBIFCH00426881 | 1997 | 4 | MÜHLEBERG | 46.92848 | 7.2745 | NMBE |
| *B. ruderatus* | GBIFCH00426872 | 1997 | 4 | BELP | 46.92859 | 7.47146 | NMBE |
| *B. ruderatus* | GBIFCH00426864 | 1998 | 4 | BELP | 46.92859 | 7.47146 | NMBE |
| *B. ruderatus* | GBIFCH00426898 | 1998 | 4 | BELP | 46.92859 | 7.47146 | NMBE |
| *B. ruderatus* | GBIFCH00426905 | 2000 | 5 | BELP | 46.92859 | 7.47146 | NMBE |
| *B. ruderatus* | GBIFCH0064042 | 2013 | 6 | Dorigny | 46.52039 | 6.55899 | PRUN |
| *B. ruderatus* | GBIFCH00499087 | 2016 | 6 | Lausanne, Maladière | 46.52087 | 6.62414 | MZL |
| *B. ruderatus* | GBIFCH00123066 | 2017 | 6 | Chancy, | 46.15456 | 5.98235 | PRUN |
| *B. ruderatus* | GBIFCH00117583 | 2018 | 6 | Gondiswil | 47.15267 | 7.86716 | MHNN |
| *B. ruderatus* | GBIFCH00139209 | 2018 | 6 | St Blaise | 47.01775 | 7.01117 | MHNN |
| *B. ruderatus* | GBIFCH00128834 | 2018 | 6 | Leuzigen | 47.19844 | 7.47162 | MHNN |
| *B. ruderatus* | GBIFCH00127953 | 2018 | 6 | Heimberg | 46.79355 | 7.60236 | MHNN |
| *B. ruderatus* | GBIFCH00135594 | 2018 | 6 | Ferreyres | 46.65479 | 6.49147 | MHNN |
| *B. ruderatus* | GBIFCH0064694 | 2018 | 6 | Chamblon | 46.74575 | 6.62074 | PRUN |
| *B. ruderatus* | GBIFCH049263 | 2019 | 6 | Trélex | 46.42674 | 6.1702 | MHNN |
| *B. ruderatus* | GBIFCH00093478 | 2019 | 6 | Bonvillars | 46.83615 | 6.68491 | MHNN |
| *B. ruderatus* | GBIFCH00135326 | 2019 | 6 | Ins | 47.01797 | 7.07694 | PRUN |
| *B. ruderatus* | GBIFCH00135325 | 2019 | 6 | Grandson, La Perraudettaz | 46.79072 | 6.62006 | PRUN |
| *B. ruderatus* | GBIFCH00123004 | 2020 | 7 | Avusy | 46.15538 | 6.04706 | MHNN |
| *B. ruderatus* | GBIFCH00123003 | 2020 | 7 | Avusy | 46.15538 | 6.04706 | MHNN |
| *B. ruderatus* | GBIFCH049660 | 2021 | 7 | Bursins | 46.42812 | 6.30028 | MHNN |
| *B. ruderatus* | GBIFCH00092929 | 2021 | 7 | Bursins | 46.42812 | 6.30028 | MHNN |
| *B. ruderatus* | GBIFCH00093241 | 2021 | 7 | Yens | 46.5193 | 6.42868 | MHNN |
| *B. ruderatus* | GBIFCH00093117 | 2021 | 7 | Daillens | 46.61034 | 6.55752 | MHNN |
| *B. ruderatus* | GBIFCH00093744 | 2021 | 7 | Orges | 46.8357 | 6.61937 | MHNN |
| *B. ruderatus* | GBIFCH00093150 | 2021 | 7 | Essertines-sur-Yverdon | 46.74575 | 6.62074 | MHNN |
| *B. ruderatus* | GBIFCH00093100 | 2021 | 7 | Daillens | 46.61034 | 6.55752 | MHNN |
| *B. ruderatus* | GBIFCH00137818 | 2021 | 7 | Bournens | 46.61034 | 6.55752 | PRUN |
| *B. ruderatus* | GBIFCH00137819 | 2021 | 7 | Oulens-sous-Echallens | 46.65531 | 6.55679 | PRUN |
| *B. ruderatus* | GBIFCH00137816 | 2021 | 7 | Essertines-sur-Yverdon | 46.74575 | 6.62074 | PRUN |
| *B. ruderatus* | GBIFCH00137814 | 2021 | 7 | Grandcour | 46.88256 | 6.94665 | PRUN |
| *B. ruderatus* | GBIFCH00137833 | 2023 | 7 | Le Landeron | 47.06295 | 7.07663 | PRAZ |
| **SPECIES** | **GBIF** | **YEAR** | **CLUSTER** | **SAMPLING LOCALITY** | **LATITUDE** | **LONGITUDE** | **MUSEUM** |
| *B. veteranus* | GBIFCH00237528 | 1941 | 1 | Lausanne, Bois de Belmont | 46.52124 | 6.67627 | MZL |
| *B. veteranus* | GBIFCH00237522 | 1941 | 1 | Lausanne, Bois de Belmont | 46.52124 | 6.67627 | MZL |
| *B. veteranus* | GBIFCH00237524 | 1941 | 1 | Vaud, Ste Catherine | 46.56647 | 6.71476 | MZL |
| *B. veteranus* | GBIFCH00237535 | 1942 | 1 | Neuchâtel, Auvernier | 46.97220 | 6.88012 | MZL |
| *B. veteranus* | GBIFCH00237489 | 1942 | 1 | Châtel s/Montsalvens, Fbg. | 46.61330 | 7.11881 | MZL |
| *B. veteranus* | GBIFCH00237537 | 1942 | 1 | Neuchâtel, Auvernier | 46.97220 | 6.88012 | MZL |
| *B. veteranus* | GBIFCH00237531 | 1942 | 1 | Neuchâtel, Auvernier | 46.97220 | 6.88012 | MZL |
| *B. veteranus* | GBIFCH00237508 | 1943 | 1 | Saint-Sulpice (VD) | 46.51129 | 6.54611 | MZL |
| *B. veteranus* | GBIFCH00237521 | 1943 | 1 | Saint-Sulpice (VD) | 46.51129 | 6.54611 | MZL |
| *B. veteranus* | GBIFCH00237548 | 1943 | 1 | Saint-Sulpice (VD) | 46.51129 | 6.54611 | MZL |
| *B. veteranus* | GBIFCH00237529 | 1943 | 1 | Cossonay | 46.60992 | 6.50531 | MZL |
| *B. veteranus* | GBIFCH00237518 | 1943 | 1 | Cheseaux-sur-Lausanne | 46.58374 | 6.61015 | MZL |
| *B. veteranus* | GBIFCH00237007 | 1944 | 1 | Neuchâtel, Auvernier | 46.80871 | 6.61978 | MZL |
| *B. veteranus* | GBIFCH00236876 | 1944 | 1 | Lausanne | 46.52865 | 6.63852 | MZL |
| *B. veteranus* | GBIFCH00236941 | 1944 | 1 | Saint-Sulpice (VD) | 46.51238 | 6.56026 | MZL |
| *B. veteranus* | GBIFCH00236954 | 1944 | 1 | Cologny | 46.21879 | 6.18480 | MZL |
| *B. veteranus* | GBIFCH00236969 | 1944 | 1 | Cologny | 46.21879 | 6.18480 | MZL |
| *B. veteranus* | GBIFCH00236873 | 1945 | 1 | Lausanne, Bois de Belmont | 46.52630 | 6.68654 | MZL |
| *B. veteranus* | GBIFCH00236936 | 1945 | 1 | Neuchâtel, Auvernier | 46.97669 | 6.87827 | MZL |
| *B. veteranus* | GBIFCH00237514 | 1945 | 1 | Vaud, St. Sulpice | 46.51129 | 6.54611 | MZL |
| *B. veteranus* | GBIFCH00237553 | 1945 | 1 | Lausanne | 46.52087 | 6.62414 | MZL |
| *B. veteranus* | GBIFCH00237527 | 1945 | 1 | Vaud, Chalet à Gobet | 46.56630 | 6.68868 | MZL |
| *B. veteranus* | GBIFCH00431836 | 1946 | 1 | Ermatingen | 47.66994 | 9.08335 | MZL |
| *B. veteranus* | GBIFCH00236871 | 1946 | 1 | Lausanne, Bois de Belmont | 46.52630 | 6.68654 | MZL |
| *B. veteranus* | GBIFCH00236957 | 1946 | 1 | Neuchâtel, Auvernier | 46.97669 | 6.87827 | MZL |
| *B. veteranus* | GBIFCH00237000 | 1946 | 1 | Neuchâtel, Auvernier | 46.97669 | 6.87827 | MZL |
| *B. veteranus* | GBIFCH00236959 | 1947 | 1 | Vaud, Entreroches | 46.66293 | 6.54650 | MZL |
| *B. veteranus* | GBIFCH00236973 | 1947 | 1 | Cologny | 46.91059 | 7.49770 | NMBE |
| *B. veteranus* | GBIFCH00271188 | 1947 | 1 | Hte Savoie, Pied du Salève | 46.84516 | 8.19249 | NMBE |
| *B. veteranus* | GBIFCH00236863 | 1948 | 1 | Vaud, St. Sulpice | 46.89257 | 7.53705 | NMBE |
| *B. veteranus* | GBIFCH00431426 | 1948 | 1 | Bern | 46.94655 | 7.35326 | NMBE |
| *B. veteranus* | GBIFCH00427018 | 1950 | 2 | Bern, Bethlehem | 46.94658 | 7.39266 | NMBE |
| *B. veteranus* | GBIFCH00427064 | 1950 | 2 | Bern | 46.94659 | 7.44520 | NMBE |
| *B. veteranus* | GBIFCH00426958 | 1950 | 2 | Bern | 46.94658 | 7.40579 | NMBE |
| *B. veteranus* | GBIFCH00431362 | 1951 | 2 | Bremgarten, Riedern | 46.95024 | 7.36650 | NMBE |
| *B. veteranus* | GBIFCH00431487 | 1951 | 2 | Belpberg | 46.94658 | 7.40580 | NMBE |
| *B. veteranus* | GBIFCH00427027 | 1951 | 2 | Köniz, Liebefeld | 46.92859 | 7.41894 | NMBE |
| *B. veteranus* | GBIFCH00426989 | 1951 | 2 | Hunzigen | 46.89257 | 7.53705 | NMBE |
| *B. veteranus* | GBIFCH00427022 | 1951 | 2 | Belp, Belpmoos | 46.91059 | 7.49770 | NMBE |
| *B. veteranus* | GBIFCH00426956 | 1952 | 2 | Bern | 46.94655 | 7.35325 | NMBE |
| *B. veteranus* | GBIFCH00427003 | 1952 | 2 | Röthenbach i.E., Schallenberg | 46.82908 | 7.79906 | NMBE |
| *B. veteranus* | GBIFCH00426995 | 1952 | 2 | Belp, Belpmoos | 46.91059 | 7.49770 | NMBE |
| *B. veteranus* | GBIFCH00426953 | 1955 | 2 | Giswil, Aaried | 46.84516 | 8.19249 | NMBE |
| *B. veteranus* | GBIFCH00427020 | 1956 | 2 | Zollikofen | 46.99156 | 7.45835 | MZL |
| *B. veteranus* | GBIFCH00236961 | 1958 | 2 | Neuchâtel, Les Ponts-de-Martel | 46.58374 | 6.61015 | MZL |
| *B. veteranus* | GBIFCH00237545 | 1958 | 2 | Neuchâtel, Les Ponts-de-Martel | 46.99839 | 6.73522 | MZL |
| *B. veteranus* | GBIFCH00427084 | 1976 | 3 | MELTINGEN | 47.37822 | 7.60414 | NMBE |
| *B. veteranus* | GBIFCH00427052 | 1978 | 3 | BUCHEGG | 47.15343 | 7.53752 | NMBE |
| *B. veteranus* | GBIFCH00427047 | 1990 | 4 | SISSELN | 47.55691 | 7.99000 | NMBE |
| *B. veteranus* | GBIFCH00427016 | 1991 | 4 | BELP | 46.92859 | 7.47146 | NMBE |
| *B. veteranus* | GBIFCH00237554 | 2010 | 5 | Ormont-Dessous | 46.43328 | 7.08085 | MZL |
| *B. veteranus* |  | 2014 | 5 | Mümliswil-Ramiswil | 47.36794 | 7.69183 | PRUN |
| *B. veteranus* | GBIFCH00118779 | 2016 | 5 | Les Ponts-de-Martel | 47.01646 | 6.74814 | MHNN |
| *B. veteranus* | GBIFCH00106334 | 2016 | 5 | Einsiedeln, Breitried | 47.09977 | 8.85476 | MHNN |
| *B. veteranus* |  | 2017 | 5 | Zürich, Reckenholz | 47.42780 | 8.52250 | ART |
| *B. veteranus* | GBIFCH00127761 | 2017 | 5 | Tourbière de la Chaux des Breuleux | 47.24284 | 7.07541 | MHNN |
| *B. veteranus* | GBIFCH00106528 | 2018 | 5 | St Imer | 47.19738 | 6.94375 | MHNN |
| *B. veteranus* | GBIFCH00132906 | 2019 | 5 | Le Chenit | 46.56165 | 6.16703 | MHNN |
| *B. veteranus* | GBIFCH00132886 | 2019 | 5 | Couvaloup de Crans | 46.42599 | 6.10517 | MHNN |
| *B. veteranus* | GBIFCH00132331 | 2019 | 5 | Le Chenit, La Burtignière | 46.56165 | 6.16703 | MHNN |
| *B. veteranus** | GBIFCH00427022 | 1951 |  | Belp, Belpmoos | 46.91059 | 7.4977 | NMBE |
| **SPECIES** | **GBIF** | **YEAR** | **CLUSTER** | **SAMPLING LOCALITY** | **LATITUDE** | **LONGITUDE** | **MUSEUM** |
| *B. pomorum* | NMB-HYMEN0000022 | 1930 | 1 | Bern, Bätterkinden | 46.21987 | 6.17503 | Bâle |
| *B. pomorum* | GBIFCH00235631 | 1933 | 1 | Suisse, Neuchâtel, Auvernier | 46.97220 | 6.88012 | MZL |
| *B. pomorum* | GBIFCH00235113 | 1934 | 1 | Vaud, La Chaux (Cossonay) | 46.61870 | 6.47905 | MZL |
| *B. pomorum* | GBIFCH00235602 | 1934 | 1 | Vaud, La Chaux (Cossonay) | 46.61870 | 6.47905 | MZL |
| *B. pomorum* | GBIFCH00235333 | 1935 | 1 | Vaud, St. Sulpice | 46.51238 | 6.56026 | MZL |
| *B. pomorum* | GBIFCH00235284 | 1936 | 1 | Lausanne, Bois de Belmont | 46.52124 | 6.67627 | MZL |
| *B. pomorum* | GBIFCH00235119 | 1936 | 1 | Vaud, St. Sulpice | 46.61870 | 6.47905 | MZL |
| *B. pomorum* | GBIFCH00235254 | 1936 | 1 | Vaud, St. Sulpice | 46.61870 | 6.47905 | MZL |
| *B. pomorum* | GBIFCH00235229 | 1936 | 1 | Vaud, Romanel-sur-Lausanne | 46.56566 | 6.59738 | MZL |
| *B. pomorum* | GBIFCH00235243 | 1936 | 1 | Vaud, St. Sulpice | 46.51238 | 6.56026 | MZL |
| *B. pomorum* | GBIFCH00235346 | 1936 | 1 | Vaud, St. Sulpice | 46.51238 | 6.56026 | MZL |
| *B. pomorum* | GBIFCH00235568 | 1936 | 1 | Vaud, St. Sulpice | 46.51238 | 6.56026 | MZL |
| *B. pomorum* | GBIFCH00235595 | 1936 | 1 | Vaud, St. Sulpice | 46.51238 | 6.56026 | MZL |
| *B. pomorum* | GBIFCH00235114 | 1939 | 1 | Vaud, Cheseaux-sur-Lausanne | 46.58374 | 6.61015 | MZL |
| *B. pomorum* | GBIFCH00235249 | 1939 | 1 | Vaud, Cheseaux-sur-Lausanne | 46.58374 | 6.61015 | MZL |
| *B. pomorum* | GBIFCH00235298 | 1939 | 1 | Vaud, Cheseaux-sur-Lausanne | 46.58374 | 6.61015 | MZL |
| *B. pomorum* | GBIFCH00235464 | 1939 | 1 | Vaud, Cheseaux-sur-Lausanne | 46.58374 | 6.61015 | MZL |
| *B. pomorum* | GBIFCH00235183 | 1940 | 2 | Vaud, Cheseaux-sur-Lausanne | 46.56566 | 6.59738 | MZL |
| *B. pomorum* | GBIFCH00235559 | 1942 | 2 | Neuchâtel, Auvernier | 46.97220 | 6.88012 | MZL |
| *B. pomorum* | GBIFCH00235273 | 1943 | 2 | Lausanne | 46.52482 | 6.62616 | MZL |
| *B. pomorum* | GBIFCH00235244 | 1943 | 2 | Cologny | 46.21987 | 6.17503 | MZL |
| *B. pomorum* | GBIFCH00235562 | 1944 | 2 | Cologny | 46.21987 | 6.17503 | MZL |
| *B. pomorum* | GBIFCH00235617 | 1944 | 2 | Cologny | 46.21987 | 6.17503 | MZL |
| *B. pomorum* | GBIFCH00235306 | 1945 | 2 | Neuchâtel, Auvernier | 46.97220 | 6.88012 | MZL |
| *B. pomorum* | GBIFCH00235286 | 1945 | 2 | Cologny | 46.21987 | 6.17503 | MZL |
| *B. pomorum* | GBIFCH00235385 | 1945 | 2 | Lausanne, Bois de Belmont | 46.52124 | 6.67627 | MZL |
| *B. pomorum* | GBIFCH00235445 | 1945 | 2 | Cologny | 46.21987 | 6.17503 | MZL |
| *B. pomorum* | GBIFCH00235570 | 1945 | 2 | Vidy | 46.52068 | 6.59808 | MZL |
| *B. pomorum* | GBIFCH00235633 | 1945 | 2 | Lausanne, Bois de Belmont | 46.52124 | 6.67627 | MZL |
| *B. pomorum* | GBIFCH00235129 | 1945 | 2 | St. Sulpice VD | 46.51238 | 6.56026 | MZL |
| *B. pomorum* | GBIFCH00235164 | 1945 | 2 | Cologny | 46.21987 | 6.17503 | MZL |
| *B. pomorum* | GBIFCH00235329 | 1946 | 2 | Neuchâtel, Auvernier | 46.97220 | 6.88012 | MZL |
| *B. pomorum* | GBIFCH00235420 | 1946 | 2 | Neuchâtel, Auvernier | 46.97220 | 6.88012 | MZL |
| *B. pomorum* | GBIFCH00235242 | 1947 | 2 | Neuchâtel, Auvernier | 46.97220 | 6.88012 | MZL |
| *B. pomorum* | GBIFCH00235339 | 1948 | 2 | Neuchâtel, Corcelles | 46.98120 | 6.88003 | MZL |
| *B. pomorum* | GBIFCH00426769 | 1949 | 2 | Bern, Liebefeld | 46.92859 | 7.41894 | NMBE |
| *B. pomorum* | GBIFCH00426767 | 1950 | 3 | Freiburg | 46.80582 | 7.15716 | NMBE |
| *B. pomorum* | GBIFCH00426739 | 1950 | 3 | Bern, Bethlehem | 46.94658 | 7.39266 | NMBE |
| *B. pomorum* | GBIFCH00426766 | 1950 | 3 | Bern, Stöckacker | 46.94658 | 7.40580 | NMBE |
| *B. pomorum* | GBIFCH00426761 | 1951 | 3 | Stöckacker BE | 46.94658 | 7.40580 | NMBE |
| *B. pomorum* | GBIFCH00426726 | 1952 | 3 | Landorf BE | 46.91959 | 7.40581 | NMBE |
| *B. pomorum* | GBIFCH00235238 | 1955 | 3 | Vaud, Bremblens | 46.54706 | 6.51944 | MZL |

**Table S2. List of primers.** For each primer used in the study the annealing temperature in Celsius and the expected size range in bp have been reported as well as the publication where they were first described.

| **PRIMER** | **ANNEALING T** | | **REFERENCE** | **SIZE RANGE (bp)** |
| --- | --- | --- | --- | --- |
| B131 | 55 °C | (Estoup et al., 1996) | | 128-154 |
| B11 | 58°C |  |  | 170-199 |
| B132 | 58°C |  |  | 128-166 |
| B100 | 65°C |  |  | 148-156 |
| B116 | 63°C |  |  | 174-177 |
| B121 | 58°C |  |  | 124-180 |
| B126 | 56°C |  |  | 158-178 |
| BT10 | 55°C | (Reber Funk et al., 2006) | | 135-159 |
| BT08 | 55°C |  |  | 142-164 |
| BT04 | 60°C |  |  | 144-196 |
| BL02 | 63 °C |  |  | 144-150 |

**Table S3. List of primers per species.** The polymorphic markers that were included in the analyses are indicated with V while with a X were reported the monomorphic primers that were excluded from the genetic analyses.

| SPECIES | B131 | BT10 | BT08 | B11 | B132 | B100 | BT04 | B116 | BL02 | B121 | B126 |
| --- | --- | --- | --- | --- | --- | --- | --- | --- | --- | --- | --- |
| *B. humilis* | V | V | V | V | V | V | X | V | X | V | V |
| *B. sylvarum* | V | V | V | V | V | X | V | V | X | V | V |
| *B. ruderatus* | V | V | V | V | V | V | V | V | V | X | V |
| *B. veteranus* | V | V | V | V | V | X | V | V | X | V | X |
| *B. pomorum* | V | V | V | V | X | X | V | V | X | V | V |

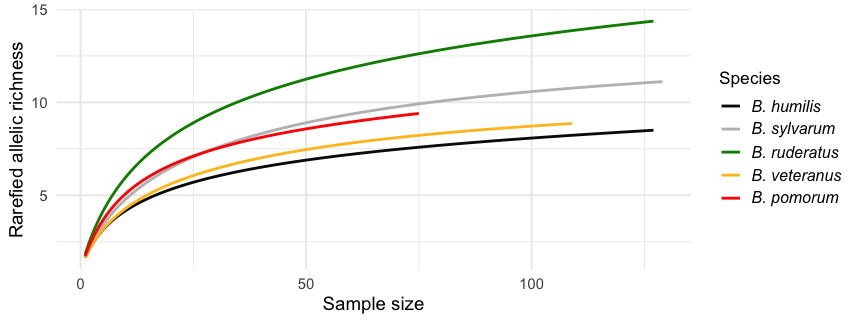

**Figure S2. Rarefaction analysis with ADZE.** Rarefaction curves with ADZE software for the number of alleles for *B. humilis, B. sylvarum, B. ruderatus, B. veteranus* and *B. pomorum*. The maximum standardized sample size was setted at 150 individuals.

**Table S4. Hardy - Weinberg equilibrium.** For each species *p-values* are reported by decade. Missing values for decades were indicated with “-”.

| **Decade** | ***B. humilis*** | ***B. sylvarum*** | ***B. ruderatus*** | ***B. veteranus*** | ***B. pomorum*** |
| --- | --- | --- | --- | --- | --- |
| **1920-1929** | 1 | 0.766 | **-** | **-** | - |
| **1930-1939** | 0.057 | < 1.78e-10 | **3.13E-07** | **-** | <6.50E-04 |
| **1940-1949** | < 5.76E-07 | 0.171 | **0.017** | **-** | < 1.78e-05 |
| **1950-1959** | 0.961 | - | **0.202** | **0.925** | 0.029 |
| **1970-1979** | - | - | **-** | **0.186** | - |
| **1980-1989** | 0.297 | - | **-** | **-** | - |
| **1990-1999** | 0.957 | - | **0.832** | **5.71E-05** | - |
| **2000-2009** | 0.07 | - | **-** | **0.903** | - |
| **2010-2019** | 1.60E-07 | < 1.41e-07 | **3.77E-08** | **0.619** | - |
| **2020-2023** | 0.469 | 0.238 | **<2.18E-04** | **<1.89E-03** | - |
| **All** | **< 1.05E-06** | **< 4.40E-17** | **< 9.24E-35** | **<6.13E-04** | **< 2.35E-07** |

**Table S5. Linkage disequilibrium.** For each species the resulting p-value (*p*) from the linkage disequilibrium inference between pair of loci **(**Locus Pair) has been reported. Significant p-values (< 0.05) have been highlighted in bold and missing locus pairs p-value are substituted with NA.

| Locus Pair | *B. HUMILIS* | *B. SYLVARUM* | *B. RUDERATUS* | *B. VETERANUS* | *B. POMORUM* |
| --- | --- | --- | --- | --- | --- |
| B100_B11 | **0.048** | NA | 0.145 | NA | NA |
| B100_B116 | 0.664 | NA | **0.000** | NA | NA |
| B100_B121 | 0.606 | NA | NA | NA | NA |
| B100_B126 | 0.516 | NA | 0.999 | NA | NA |
| B100_B131 | 0.411 | NA | NA | NA | NA |
| B100_B132 | 0.547 | 0.271 | **0.000** | NA | NA |
| B100_BL02 | NA | NA | **0.000** | NA | NA |
| B100_BT04 | NA | NA | 0.210 | NA | NA |
| B100_BT08 | 0.135 | NA | **0.000** | NA | NA |
| B100_BT10 | 0.105 | NA | 0.748 | NA | NA |
| B116_B121 | 0.158 | 0.933 | NA | 1.000 | 0.156 |
| B116_B126 | 0.234 | 0.904 | 0.736 | NA | 0.725 |
| B116_B131 | **0.033** | 0.182 | NA | 0.729 | 0.869 |
| B116_B132 | 0.342 | 0.721 | **0.004** | 0.987 | **0.004** |
| B116_BL02 | NA | NA | **0.000** | NA | NA |
| B116_BT04 | NA | NA | **0.013** | 0.219 | 0.414 |
| B116_BT08 | 0.192 | 0.422 | **0.000** | 0.457 | 0.194 |
| B116_BT10 | 0.627 | 0.124 | 0.968 | 0.640 | 0.879 |
| B11_B116 | 0.388 | 0.783 | **0.000** | 0.677 | 0.993 |
| B11_B121 | 0.268 | 0.945 | NA | 0.813 | 1.000 |
| B11_B126 | 0.734 | 0.136 | 0.109 | NA | 1.000 |
| B11_B131 | 0.931 | 0.232 | NA | 0.788 | 0.649 |
| B11_B132 | 0.830 | 0.962 | **0.000** | 0.985 | **0.000** |
| B11_BL02 | NA | NA | **0.000** | NA | NA |
| B11_BT04 | NA | NA | 0.752 | 0.982 | 0.595 |
| B11_BT08 | 0.682 | **0.025** | **0.001** | **0.043** | 0.892 |
| B11_BT10 | 0.730 | 0.565 | **0.048** | 0.321 | 1.000 |
| B121_B126 | 0.125 | 0.643 | NA | NA | 1.000 |
| B121_B131 | 0.224 | 0.674 | NA | 0.614 | 0.641 |
| B121_B132 | **0.001** | NA | NA | NA | NA |
| B121_BT04 | NA | NA | NA | 0.726 | 0.702 |
| B121_BT08 | 0.231 | 1.000 | NA | 0.860 | 0.296 |
| B121_BT10 | 0.450 | 0.113 | NA | 0.390 | 1.000 |
| B126_B131 | 0.222 | 0.066 | NA | NA | 0.165 |
| B126_B132 | **0.006** | **0.022** | 0.090 | NA | 0.769 |
| B126_BT04 | NA | NA | 0.691 | NA | NA |
| B126_BT08 | 0.388 | 0.558 | NA | NA | 0.315 |
| B126_BT10 | 0.220 | 0.949 | **0.032** | NA | 1.000 |
| B131_B132 | **0.025** | 0.959 | NA | 0.129 | **0.003** |
| B131_BT04 | NA | NA | NA | 0.136 | 0.135 |
| B131_BT08 | 0.721 | 0.828 | NA | 0.092 | 0.485 |
| B131_BT10 | 0.862 | 0.916 | NA | 0.452 | **0.000** |
| B132_BL02 | NA | NA | **0.000** | NA | NA |
| B132_BT04 | NA | NA | **0.005** | **0.019** | NA |
| B132_BT08 | 0.586 | 0.429 | **0.000** | 0.717 | **0.000** |
| B132_BT10 | 0.621 | 0.544 | **0.000** | 0.382 | **0.000** |
| BL02_BT04 | NA | NA | 0.084 | NA | NA |
| BL02_BT10 | NA | NA | **0.000** | NA | NA |
| BT04_BT08 | NA | NA | **0.036** | 0.398 | 0.917 |
| BT04_BT10 | NA | NA | 0.567 | 0.353 | **0.038** |
| BT08_BT10 | 0.442 | **0.000** | 0.666 | 0.821 | 0.697 |

**Table S6. Observed heterozygosity per locus in each species.** For each species the observed heterozygosity for each locus. Missing locus observed heterozygosity are substituted with NA and observed heterozygosity values lower than 0.5 are in bold.

| **Locus** | ***B. humilis*** | ***B. sylvarum*** | ***B. ruderatus*** | ***B. veteranus*** | ***B. pomorum*** |
| --- | --- | --- | --- | --- | --- |
| B131 | 0.710 | **0.458** | 0.551 | 0.714 | **0.254** |
| BT08 | 0.779 | 0.747 | 0.539 | 0.738 | 0.559 |
| B11 | **0.297** | **0.468** | **0.486** | 0.805 | **0.481** |
| BT10 | 0.737 | 0.539 | 0.551 | 0.838 | 0.661 |
| B116 | **0.268** | **0.463** | **0.299** | **0.366** | 0.519 |
| B126 | **0.368** | **0.357** | 0.711 | 0.762 | NA |
| B100 | **0.000** | NA | **0.397** | NA | NA |
| B121 | 0.594 | 0.677 | NA | 0.667 | 0.589 |
| B132 | 0.783 | 0.609 | **0.446** | **0.123** | **0.123** |
| BT04 | NA | 0.568 | 0.675 | 0.500 | 0.685 |
| BL02 | NA | NA | **0.360** | NA | NA |

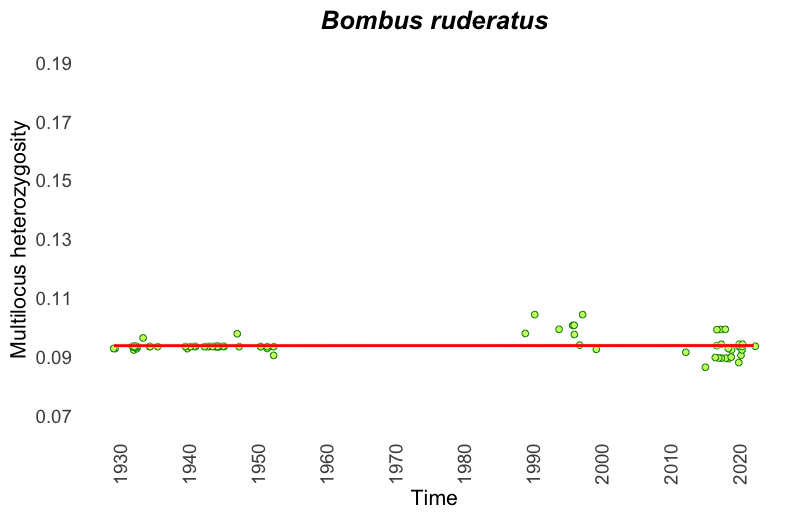

**Figure S3. Multilocus heterozygosity of *B. ruderatus* from 1930-2023.** Linear correlation between multilocus heterozygosity of *B. ruderatus* time have been represented. A 95% of confidence interval has been also shown towards the intercept line.

**Table S7. Robust linear regression model for multilocus heterozygosity.** Results of robust linear regression models testing for temporal trends in genetic diversity for each *Bombus* species. The response variable represents multilocus heterozygosity, and the predictor is collection year. Results with *p*-value < 0.05 were marked with an asterisk (*). Adjusted R² indicates the proportion of variance explained by the model, adjusted for the number of predictors.

| ***BOMBUS HUMILIS*** | | | | |
| --- | --- | --- | --- | --- |
| **Predictor** | **Estimate** | **Std. Error** | **t-value** | **p-value** |
| Intercept | 0.226 | 3.81x10^-02^ | 5.934 | 4.18x10^-08^ |
| Year | -4.64x10^-05^ | 1.94x10^-05^ | -2.392 | 1.86x10^-02^* |
| Adjusted R² | 0.052 |  |  |  |
| **BOMBUS *SYLVARUM*** | | | | |
| **Predictor** | **Estimate** | **Std. Error** | **t-value** | **p-value** |
| Intercept | –0.030 | 0.08 | –0.368 | 0.714 |
| Year | 0.0001 | 0.000 | 1.744 | 0.085 |
| Adjusted R² | 0.045 |  |  |  |
| ***BOMBUS RUDERATUS* (1929-2023)** | | | | |
| **Predictor** | **Estimate** | **Std. Error** | **t-value** | **p-value** |
| Intercept | 9.28x10^-02^ | 1.62x10^-02^ | 5.745 | 1.65x10^-07^ |
| Year | 4.85x10^-07^ | 8.33x10^-06^ | 0.058 | 0.954 |
| Adjusted R² | -0.013 |  |  |  |
| **COEFFICIENTS *B. RUDERATUS* (1929-1960)** | | | | |
| **Predictor** | **Estimate** | **Std. Error** | **t-value** | **p-value** |
| Intercept | 3.55x10^-02^ | 0.15 | 0.230 | 0.819 |
| Year | 6.52x10^-05^ | 7.96x10^-05^ | 0.819 | 0.417 |
| Adjusted R² | -5.95x10^-05^ |  |  |  |
| **COEFFICIENTS *B. RUDERATUS* (1990-2023)** | | | | |
| **Predictor** | **Estimate** | **Std. Error** | **t-value** | **p-value** |
| Intercept | 0.994 | 0.49 | 2.03 | 5.04x10^-02^ |
| Year | -4.13x10^-04^ | 2.43x10^-04^ | -1.70 | 9.82x10^-02^* |
| Adjusted R² | 0.112 |  |  |  |
| **BOMBUS *VETERANUS*** | | | | |
| **Predictor** | **Estimate** | **Std. Error** | **t-value** | **p-value** |
| Intercept | 2.45x10^-02^ | 7.49x10^-02^ | 0.327 | 0.745 |
| Year | 6.02x10^-05^ | 3.85x10^-05^ | 1.564 | 0.123 |
| Adjusted R² | 0.019 |  |  |  |
| **BOMBUS *POMORUM*** | | | | |
| **Predictor** | **Estimate** | **Std. Error** | **t-value** | **p-value** |
| Intercept | -0.131 | 0.113 | -1.159 | 0.253 |
| Year | 0.0001 | 5.83x10^-05^ | 2.249 | 0.030* |
| Adjusted R² | 0.131 |  |  |  |

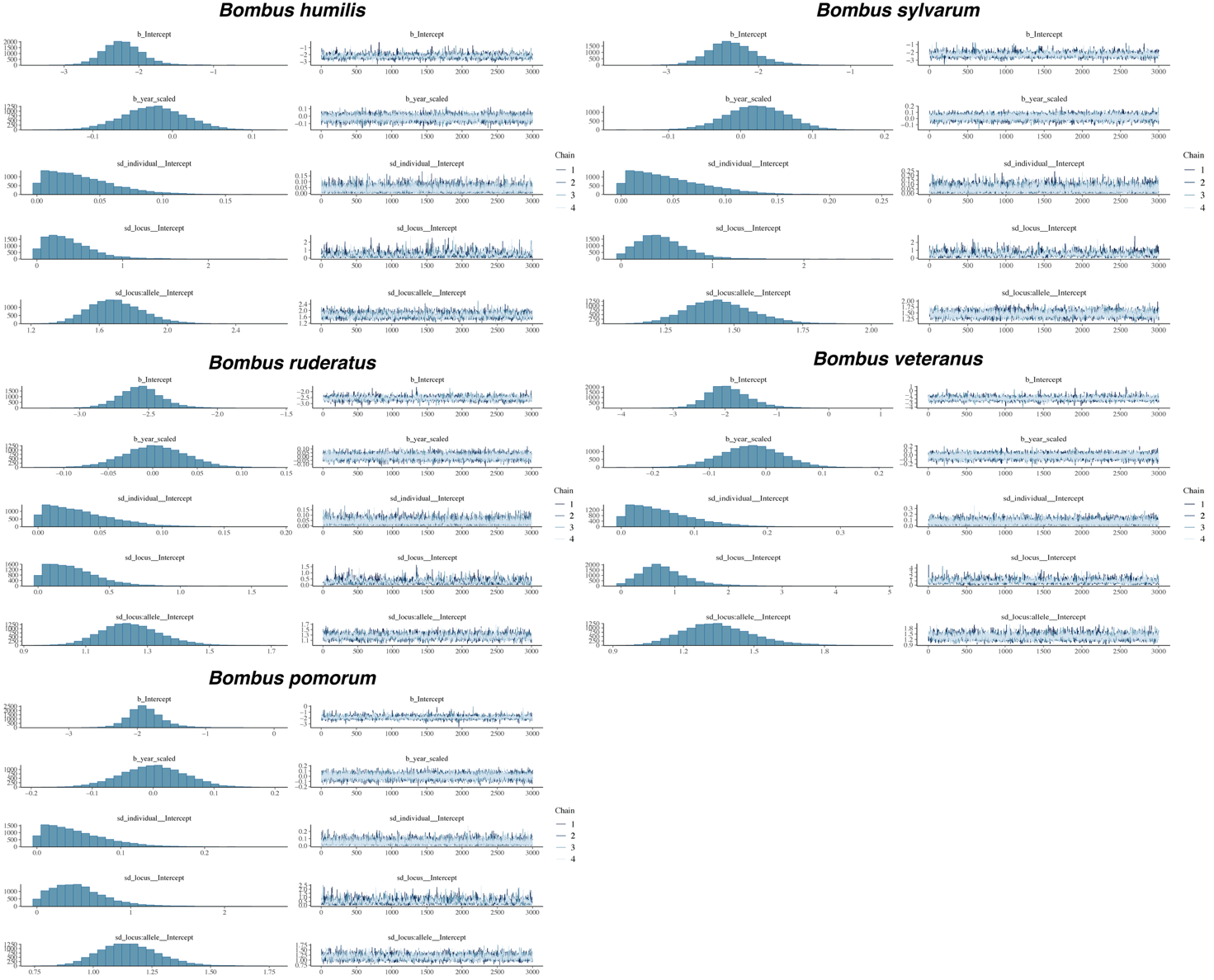

**Figure S4. Diagnostic plots Bayesian analysis of allelic richness for five *Bombus* species.** Posterior distributions and model diagnostics for the Bayesian logistic mixed-effects model predicting the presence of alleles in each species as a function of scaled year (year_scaled), with random intercepts for allele identity. The plots show the posterior distributions of the fixed effects (intercept and slope), the standard deviation of the random effect (allele), and model diagnostics including Rhat (convergence indicator, with values of 1 indicating good convergence) and effective sample sizes (Bulk_ESS and Tail_ESS). Vertical lines represent 95% credible intervals.

**Table S8. Summary of Bayesian logistic regression model results assessing the effect of scaled year on allele presence for five *Bombus* species.** For each species, models were fitted with a logit link and included scaled year as a fixed effect, with random intercepts for individual, locus, and allele nested within locus (locus:allele). Fixed effects included the model intercept (baseline log-odds of allele presence when scaled year = 0) and the effect of scaled year (temporal trend in log-odds of allele presence) while random-effect variance components were expressed as standard deviations (sd) on the logit scale. For all parameters, we report posterior means (Estimate), posterior standard errors (Est.Error), and 95% credible intervals. Convergence was assessed using the Gelman–Rubin diagnostic (R̂) and bulk effective sample sizes (Bulk ESS).

| **Species** | **Parameter** | **Estimate** | **Est.Error** | **95% CI** | **R̂** | **Bulk ESS** |
| --- | --- | --- | --- | --- | --- | --- |
| ***Bombus humilis*** | Intercept | -2.21 | 0.28 | -2.71– -1.63 | 1 | 2529 |
|  | year_scaled | -0.02 | 0.04 | -0.10–0.05 | 1 | 25727 |
|  | sd_individual | 0.04 | 0.03 | 0.00–0.10 | 1 | 9435 |
|  | sd_locus | 0.39 | 0.3 | 0.02–1.15 | 1.01 | 1243 |
|  | sd_locus:allele | 1.7 | 0.16 | 1.42–2.03 | 1 | 3112 |
| ***Bombus sylvarum*** | Intercept | -2.28 | 0.27 | -2.76– -1.70 | 1 | 2117 |
|  | year_scaled | 0.02 | 0.04 | -0.06–0.10 | 1 | 17654 |
|  | sd_individual | 0.05 | 0.04 | 0.00–0.13 | 1 | 7003 |
|  | sd_locus | 0.46 | 0.28 | 0.04–1.11 | 1 | 1173 |
|  | sd_locus:allele | 1.45 | 0.12 | 1.23–1.70 | 1 | 3242 |
| ***Bombus ruderatus*** | Intercept | -2.56 | 0.16 | -2.86– -2.22 | 1 | 2527 |
|  | year_scaled | 0 | 0.03 | -0.07–0.07 | 1 | 18046 |
|  | sd_individual | 0.03 | 0.03 | 0.00–0.10 | 1 | 7589 |
|  | sd_locus | 0.25 | 0.19 | 0.01–0.69 | 1.01 | 651 |
|  | sd_locus:allele | 1.25 | 0.1 | 1.07–1.45 | 1 | 2454 |
| ***Bombus veteranus*** | Intercept | -1.92 | 0.42 | -2.67– -0.99 | 1 | 3447 |
|  | year_scaled | -0.03 | 0.05 | -0.13–0.08 | 1 | 19646 |
|  | sd_individual | 0.06 | 0.04 | 0.00–0.16 | 1 | 7191 |
|  | sd_locus | 0.82 | 0.46 | 0.10–1.92 | 1 | 1545 |
|  | sd_locus:allele | 1.35 | 0.15 | 1.09–1.68 | 1 | 3163 |
| ***Bombus pomorum*** | Intercept | -1.88 | 0.26 | -2.34– -1.31 | 1 | 3429 |
|  | year_scaled | 0 | 0.05 | -0.10–0.10 | 1 | 18538 |
|  | sd_individual | 0.05 | 0.04 | 0.00–0.13 | 1 | 8352 |
|  | sd_locus | 0.45 | 0.3 | 0.03–1.17 | 1 | 1305 |
|  | sd_locus:allele | 1.15 | 0.13 | 0.93–1.43 | 1 | 3362 |

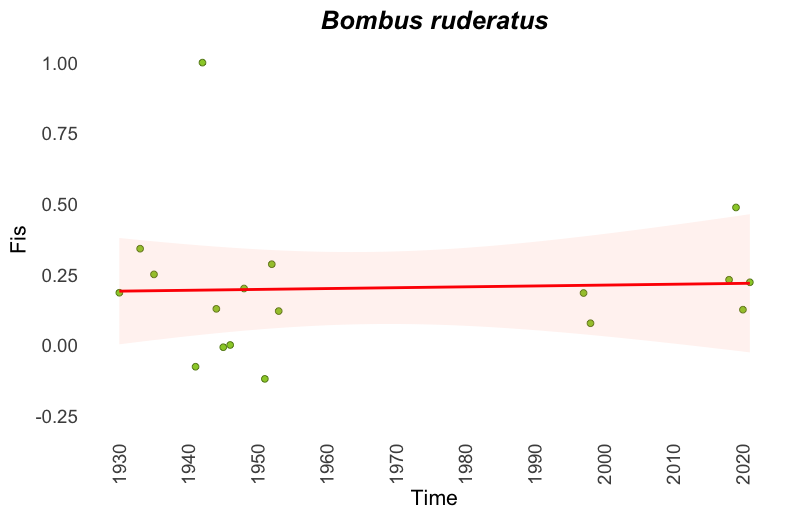

**Figure S5. Inbreeding coefficient of *B. ruderatus*.** Linear correlation between inbreeding coefficient (Fis) of *B. ruderatus* and time have been represented. A 95% of confidence interval has been also shown towards the intercept line.

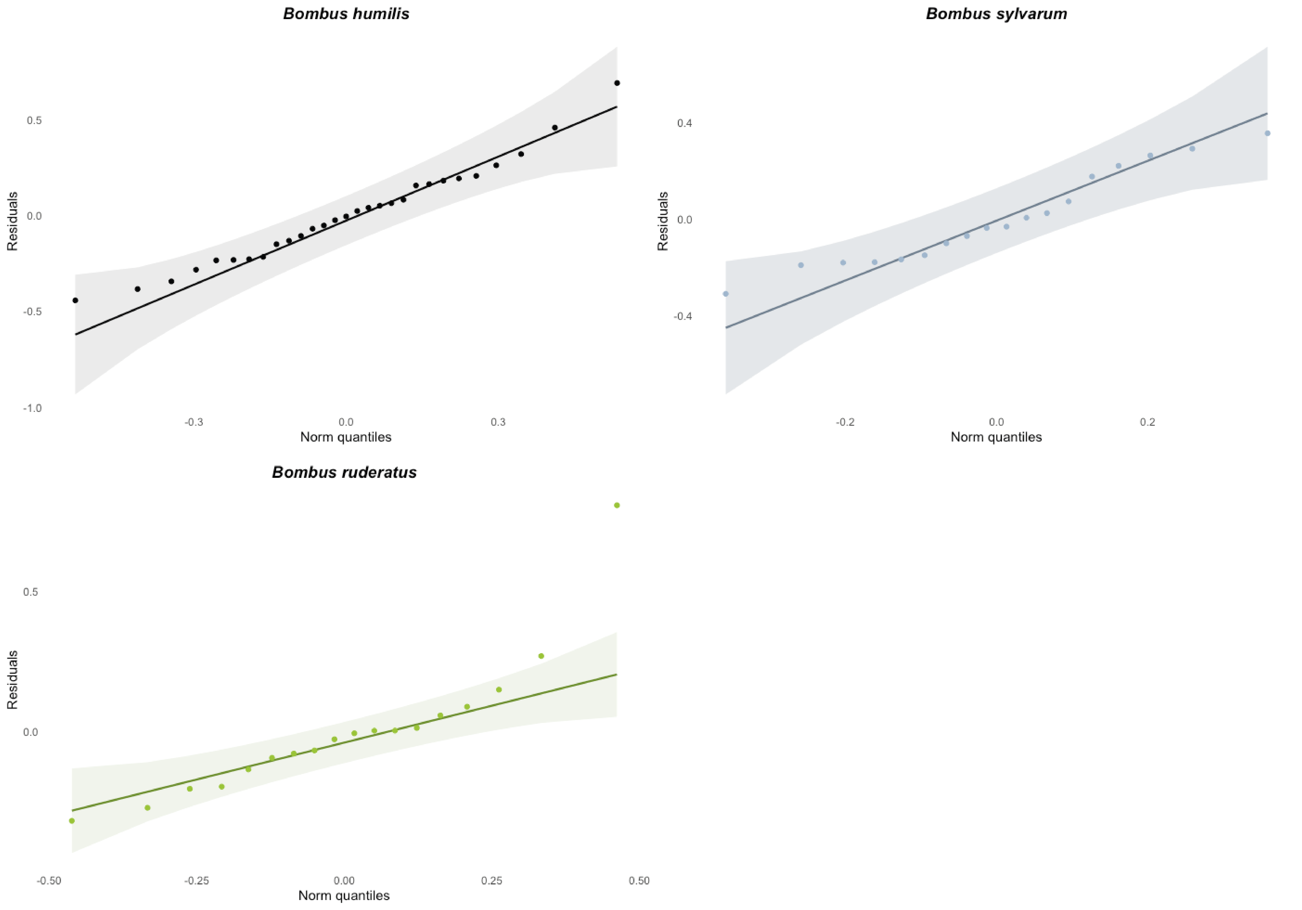

**Figure S6. Quantile-Quantile plot of the residuals from the linear model estimating inbreeding coefficient (Fis) over time.** Each plot illustrates the distribution of residuals compared to a theoretical normal distribution. The QQ line represents the expected normal distribution, while the shaded area indicates the 95% confidence interval.

**Table S9. Linear regression model of inbreeding coefficient.** Results of linear regression models testing for temporal trends in inbreeding for each *Bombus* species. The response variable represents the inbreeding coefficient (Fis), and the predictor is collection year. Results with *p*-value < 0.05 were marked with an asterisk (*). Adjusted R² indicates the proportion of variance explained by the model, adjusted for the number of predictors.

| ***BOMBUS HUMILIS*** | | | | |
| --- | --- | --- | --- | --- |
|  | **Estimate** | **Std. Error** | **t-value** | **p-value** |
| **Intercept** | -3.898 | 2.699 | -1.445 | 0.160 |
| **Time** | 0.002 | 0.001 | 1.495 | 0.147 |
| **Adjusted R2** | 0.042 |  |  |  |
| ***BOMBUS SYLVARUM*** | | | | |
|  | **Estimate** | **Std. Error** | **t-value** | **p-value** |
| **Intercept** | 1.559 | 2.427 | 0.643 | 0.530 |
| **Time** | -0.001 | 0.001 | -0.551 | 0.589 |
| **Adjusted R2** | -0.043 |  |  |  |
| ***BOMBUS RUDERATUS* (1930-2023)** | | | | |
|  | **Estimate** | **Std. Error** | **t-value** | **p-value** |
| **Intercept** | -0.400 | 3.537 | -0.113 | 0.911 |
| **Time** | 0.0003 | 0.002 | 0.170 | 0.867 |
| **Adjusted R2** | -0.061 |  |  |  |
| ***BOMBUS RUDERATUS* (1930-1960)** | | | | |
|  | **Estimate** | **Std. Error** | **t-value** | **p-value** |
| **Intercept** | 17.423 | 23.329 | 0.747 | 0.472 |
| **Time** | -0.009 | 0.012 | -0.739 | 0.477 |
| **Adjusted R2** | -0.043 |  |  |  |

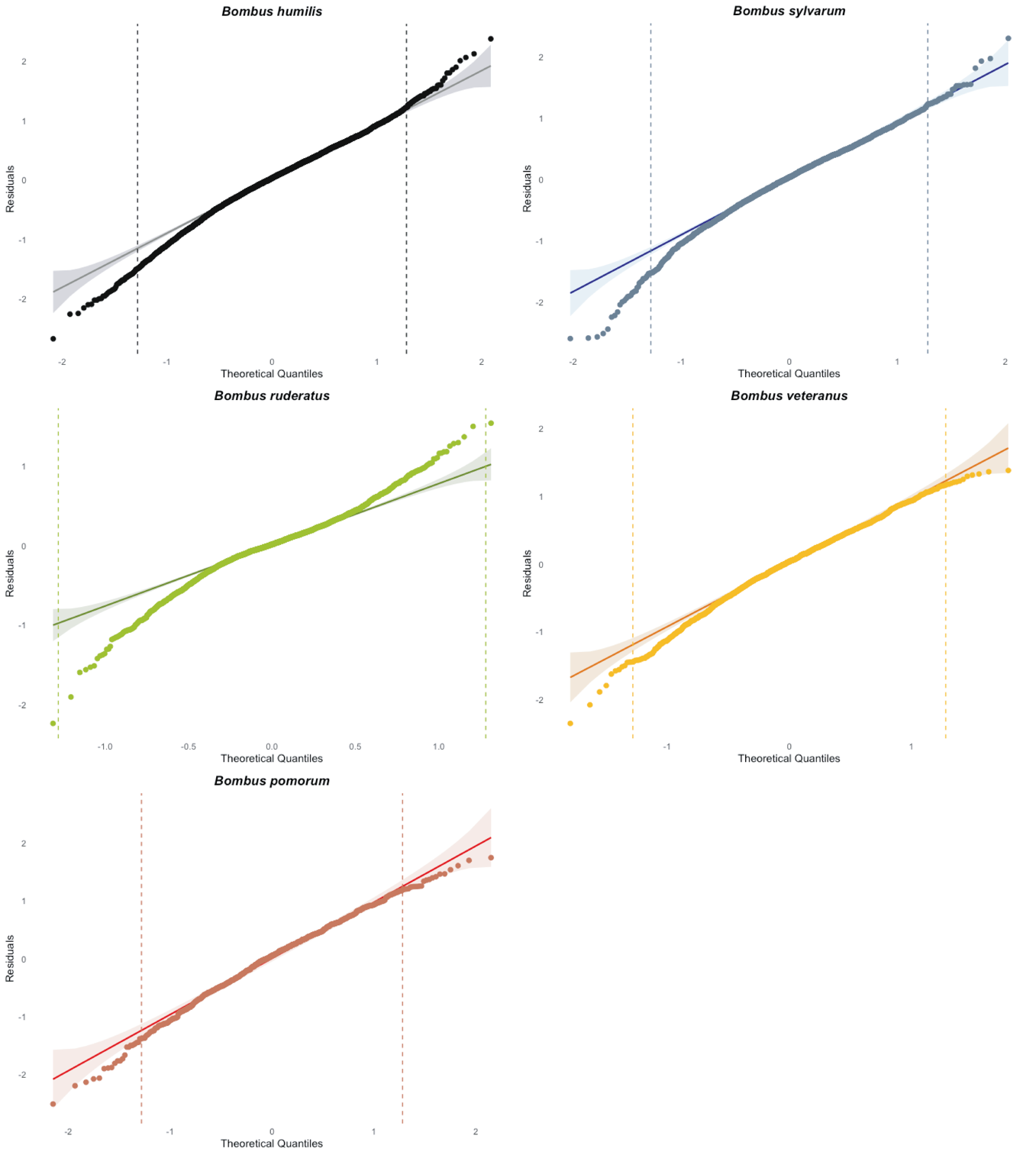

**Figure S7. QQ plot residuals of the linear model between genetic distances and geographical and time distances.** The residuals distribution is represented through a QQ plot where QQ line is represented in red and the 95% confidence interval is defined by a shaded area along the QQ line. The central quantile range of 10%-90% (z = ± 1.28) is represented by dashed lines in the QQ plot to visualize the position of 80% of the residuals.

**Table S10. Correlation analysis between temporal and spatial distance and PC1 and PC2.** Results of correlation analyses between temporal and spatial distance and the first two principal component axes (PC1 and PC2) for *B. humilis*, *B. sylvarum*, *B. ruderatus*, *B. veteranus* and *B. pomorum*. The correlation coefficient (*r*), t-statistic (*t*), and associated p-value (*p*) were reported.

| **Species** | **Explanatory variable** | **PCA axis** | ***r*** | ***t*** | ***p*** |
| --- | --- | --- | --- | --- | --- |
| *B. humilis* | Temporal distance | PC1 | –0.279 | –2.94 | 0.004 |
|  | Temporal distance | PC2 | –0.085 | –0.86 | 0.392 |
|  | Spatial distance | PC1 | 0.102 | 1.04 | 0.302 |
|  | Spatial distance | PC2 | 0.117 | 1.19 | 0.237 |
| *B. sylvarum* | Temporal distance | PC1 | 0.148 | 1.32 | 0.19 |
|  | Temporal distance | PC2 | –0.149 | –1.33 | 0.187 |
|  | Spatial distance | PC1 | –0.179 | -1.61 | 0.111 |
|  | Spatial distance | PC2 | –0.194 | –1.74 | 0.085 |
| *B. ruderatus* | Temporal distance | PC1 | 0.796 | 11.69 | < 2.2x10^-16^ |
|  | Temporal distance | PC2 | –0.250 | –2.30 | 0.024 |
|  | Spatial distance | PC1 | 0.065 | 0.58 | 0.564 |
|  | Spatial distance | PC2 | 0.018 | 0.16 | 0.872 |
| *B. veteranus* | Temporal distance | PC1 | –0.542 | –4.91 | 7.86x10^-16^ |
|  | Temporal distance | PC2 | 0.439 | 3.72 | 0.0005 |
|  | Spatial distance | PC1 | –0.289 | –2.30 | 0.025 |
|  | Spatial distance | PC2 | 0.15 | 1.16 | 0.252 |
| *B. pomorum* | Temporal distance | PC1 | 0.117 | 0.74 | 0.461 |
|  | Temporal distance | PC2 | –0.082 | –0.52 | 0.604 |
|  | Spatial distance | PC1 | 0.012 | 0.07 | 0.941 |
|  | Spatial distance | PC2 | –0.024 | –0.15 | 0.882 |

**Table S11. Redundancy analysis (RDA).** Results of redundancy analysis and analysis of variance (ANOVA) testing the effects of temporal and spatial variables on genetic variation in *B. humilis*, *B. sylvarum*, *B. ruderatus*, *B. veteranus* and *B. pomorum*. Reported are the F-statistics (F) and associated p-values (*p*) for the full RDA model (time + space), partial RDAs testing time and space separately, and ANOVA tests for individual predictors (year, longitude, and latitude).

| **Species** | **Analysis** | **Variable** | **F** | ***p*** |
| --- | --- | --- | --- | --- |
| *B. humilis* | RDA full | time + space | 4.48 | 0.001 |
|  | RDA partial | time | 10.86 | 0.001 |
|  | RDA partial | space | 1.01 | 0.429 |
|  | anova | year | 11.43 | 0.001 |
|  | anova | longitude | 1.14 | 0.265 |
|  | anova | latitude | 0.87 | 0.62 |
| *B. sylvarum* | RDA full | time + space | 1.12 | 0.186 |
|  | RDA partial | time | 2.08 | 0.001 |
|  | RDA partial | space | 0.61 | 0.997 |
|  | anova | year | 2.13 | 0.001 |
|  | anova | longitude | 0.81 | 0.796 |
|  | anova | latitude | 0.42 | 0.416 |
| *B. ruderatus* | RDA full | time + space | 9.28 | 0.001 |
|  | RDA partial | time | 24.8 | 0.001 |
|  | RDA partial | space | 1.54 | 0.001 |
|  | anova | year | 24.77 | 0.001 |
|  | anova | longitude | 1.62 | 0.07 |
|  | anova | latitude | 1.46 | 0.114 |
| *B. veteranus* | RDA full | time + space | 2.62 | 0.001 |
|  | RDA partial | time | 3.79 | 0.001 |
|  | RDA partial | space | 1.64 | 0.005 |
|  | anova | year | 4.58 | 0.001 |
|  | anova | longitude | 1.56 | 0.035 |
|  | anova | latitude | 1.71 | 0.017 |
| *B. pomorum* | RDA full | time + space | 1.09 | 0.221 |
|  | RDA partial | time | 1.32 | 0.101 |
|  | RDA partial | space | 1.21 | 0.092 |
|  | anova | year | 0.86 | 0.748 |
|  | anova | longitude | 1.59 | 0.009 |
|  | anova | latitude | 0.83 | 0.769 |

**Table S12. Linear mixed model of spatial and temporal effects on genetic differentiation.** Results of linear mixed-effects models examining the influence of geographic distance, temporal distance, and their interaction on genetic differentiation in five *Bombus* species. Fixed effects include intercept, geographic distance (geo), temporal distance (time), and geo × time interaction. Random intercepts were included for each population (id1 and id2). Both marginal and conditional R² values were reported. Significant p-values (p ≤ 0.05) are marked with an asterisk (*), and highly significant results (p ≤ 0.001) with two asterisks (**).

| ***BOMBUS HUMILIS*** | | | | | | |
| --- | --- | --- | --- | --- | --- | --- |
| **Fixed effects:** | **Estimate** | **Std. Error** | **df** | | **t-value** | **p-value** |
| Intercept | -0.234 | 8.11x10^-02^ | | 1.65x10^2^ | -2.88 | 0.005** |
| Geographic distance (geo) | 1.21x10^-03^ | 9.89x10^-03^ | | 5.28x10^3^ | 0.122 | 0.903 |
| Temporal distance (time) | 0.228 | 2.49x10^-02^ | | 9.71x10^2^ | 9.187 | <2x10^-16^** |
| geo * time interaction | -1.39x10^-02^ | 8.66x10^-03^ | | 5.22x10^3^ | -1.6 | 0.110 |
| Marginal R^2^ | 0.054 |  | |  |  |  |
| Conditional R^2^ | 0.668 |  | |  |  |  |
| **Random Effects:** | **Variance** | **Std. Dev.** | |  |  |  |
| id1 (intercept) | 0.407 | 0.638 | |  |  |  |
| id2 (intercept) | 0.192 | 0.438 | |  |  |  |
| Residual | 0.324 | 0.569 | |  |  |  |
| ***BOMBUS SYLVARUM*** | | | | | | |
| **Fixed effects** | **Estimate** | **Std. Error** | | **df** | **t-value** | **p-value** |
| Intercept | -0.052 | 0.099 | | 146.872 | -0.525 | 0.600 |
| Geographic distance (geo) | -0.010 | 0.013 | | 3085.942 | -0.749 | 0.454 |
| Temporal distance (time) | 0.087 | 0.043 | | 225.257 | 2.053 | 0.041* |
| geo * time interaction | 0.005 | 0.012 | | 3054.062 | 0.394 | 0.694 |
| Marginal R^2^ | 0.008 |  | |  |  |  |
| Conditional R^2^ | 0.656 |  | |  |  |  |
| **Random Effects** | **Variance** | **Std. Dev.** | |  |  |  |
| id1 (intercept) | 0.373 | 0.611 | |  |  |  |
| id2 (intercept) | 0.253 | 0.503 | |  |  |  |
| Residual | 0.333 | 0.577 | |  |  |  |
| ***BOMBUS RUDERATUS*** | | | | | | |
| **Fixed effects** | **Estimate** | **Std. Error** | | **df** | **t-value** | **p-value** |
| Intercept | -0.177 | 6.97x10^-02^ | | 1.45x10^2^ | -2.54 | 0.012* |
| Geographic distance (geo) | 5.52x10^-03^ | 8.00x10^-03^ | | 3.16x10^3^ | 0.69 | 0.4902 |
| Temporal distance (time) | 0.606 | 1.26x10^-02^ | | 3.13x10^3^ | 48.001 | <2x10^-16^** |
| geo * time interaction | 7.73x10^-03^ | 7.35x10^-03^ | | 3.12x10^3^ | 1.052 | 0.293 |
| Marginal R^2^ | 0.419 |  | |  |  |  |
| Conditional R^2^ | 0.841 |  | |  |  |  |
| **Random Effects** | **Variance** | **Std. Dev.** | |  |  |  |
| id1 (intercept) | 0.222 | 0.472 | |  |  |  |
| id2 (intercept) | 0.146 | 0.382 | |  |  |  |
| Residual | 0.139 | 0.373 | |  |  |  |
| ***BOMBUS VETERANUS*** | | | | | | |
| **Fixed effects** | **Estimate** | **Std. Error** | | **df** | **t-value** | **p-value** |
| Intercept | -0.1284 | 0.108 | | 106.015 | -1.194 | 0.235 |
| Geographic distance (geo) | 5.31x10^-02^ | 2.05x10^-02^ | | 1.75x10^3^ | 2.597 | 0.009** |
| Temporal distance (time) | 0.151 | 4.14x10^-02^ | | 8.50x10^2^ | 3.647 | 2.81x10^-04^** |
| geo * time interaction | -2.76x10^-02^ | 1.64x10^-02^ | | 1.70x10^3^ | -1.686 | 0.092 |
| Marginal R^2^ | 0.032 |  | |  |  |  |
| Conditional R^2^ | 0.688 |  | |  |  |  |
| **Random Effects** | **Variance** | **Std. Dev.** | |  |  |  |
| id1 (intercept) | 0.357 | 0.598 | |  |  |  |
| id2 (intercept) | 0.252 | 0.502 | |  |  |  |
| Residual | 0.289 | 0.538 | |  |  |  |
| ***BOMBUS POMORUM*** | | | | | | |
| **Fixed effects** | **Estimate** | **Std. Error** | | **df** | **t-value** | **p-value** |
| Intercept | -8.37x10^-04^ | 0.123 | | 71.3 | -0.007 | 0.995 |
| Geographic distance (geo) | 4.07x10^-02^ | 3.19x10^-02^ | | 8.49x10^2^ | 1.276 | 0.202 |
| Temporal distance (time) | -1.14x10^-02^ | 3.03x10^-02^ | | 8.43x10^2^ | -0.377 | 0.706 |
| geo * time interaction | -5.95x10^-02^ | 2.6610^-02^ | | 8.36x10^2^ | -2.237 | 0.026* |
| Marginal R^2^ | 0.006 |  | |  |  |  |
| Conditional R^2^ | 0.535 |  | |  |  |  |
| **Random Effects** | **Variance** | **Std. Dev.** | |  |  |  |
| id1 (intercept) | 0.294 | 0.542 | |  |  |  |
| id2 (intercept) | 0.251 | 0.501 | |  |  |  |
| Residual | 0.479 | 0.692 | |  |  |  |
